# Genomic architecture drives population structuring in Amazonian birds

**DOI:** 10.1101/2021.12.01.470789

**Authors:** Gregory Thom, Lucas Rocha Moreira, Romina Batista, Marcelo Gehara, Alexandre Aleixo, Brian Tilston Smith

## Abstract

Large rivers are ubiquitously invoked to explain the distributional limits and speciation of the Amazon Basin’s mega-diversity. However, inferences on the spatial and temporal origins of Amazonian species have narrowly focused on evolutionary neutral models, ignoring the potential role of natural selection and intrinsic genomic processes known to produce heterogeneity in differentiation across the genome. To test how genomic architecture impacts our ability to reconstruct patterns of spatial diversification across multiple taxa, we sequenced whole genomes for populations of bird species that co-occur in southeastern Amazonian. We found that phylogenetic relationships within species and demographic parameters varied across the genome in predictable ways. Genetic diversity was positively associated with recombination rate and negatively associated with the species tree topology weight. Gene flow was less pervasive in regions of low recombination, making these windows more likely to retain patterns of population structuring that matched the species tree. We further found that approximately a third of the genome showed evidence of selective sweeps and linked selection skewing genome-wide estimates of effective population sizes and gene flow between populations towards lower values. In sum, we showed that the effects of intrinsic genomic characteristics and selection can be disentangled from the neutral processes to elucidate how speciation hypotheses and biogeographic patterns are sensitive to genomic architecture.

## Introduction

Across the Amazon Basin, large rivers delimit the distribution of hundreds of rainforest taxa (Cracraft 1985; Bates et al. 1998; da Silva et al. 2005). The spatial patterns that underlie these distributions have been central for understanding how diversity originates in the hyperdiverse Neotropics (Silva et al. 2019; Haffer 2008, 1969; Ribas et al. 2012; Smith et al. 2014). The species isolated by large rivers show complex and highly variable relationships that span millions of years, with limited congruence in spatial patterns of diversification and historical demography (Smith et al. 2014; Silva et al. 2019). Reduced genomic approaches have revealed that factors such as gene flow may hinder inferences on the origins of species distributed across Amazonian rivers (Weir et al. 2015; Barrera-Guzmán et al. 2018; Berv et al. 2021; Ferreira et al. 2018; Luna et al. 2021; Musher et al. 2021; Del-Rio et al. 2021). In addition to gene flow, intrinsic (e.g., recombination rate) and extrinsic (e.g., selection) processes that influence the landscape of genomic diversity and differentiation may further obfuscate biogeographic inferences by affecting the estimation of phylogenetic and demographic parameters (Bravo et al. 2021; Harvey et al. 2019). Elucidating the relationships between the processes driving genomic evolution may yield more accurate inferences on the spatial and temporal history of species, providing increased resolution into the hotly debated origins of Amazonian biodiversity.

The genomic landscape of genetic diversity is ubiquitous across taxonomic groups indicating that evolutionary signal is dependent on which portions of the genome are examined (Martin et al. 2019; Li et al. 2019; Manthey et al. 2021; Delmore et al. 2018; Johri et al.). Components of genomic architecture, such as chromosome inheritance, meiotic recombination, the density of targets of selection, biased gene conversion, and mutation rate operate simultaneously and heterogeneously across the genome, resulting in highly variable levels of genetic diversity and divergence at both intra- and interspecific scales (Martin et al. 2019; Edelman et al. 2019; Garrigan et al. 2012; Fontaine et al. 2015; Smith et al. 2018; Meunier and Duret 2004; Cruickshank and Hahn 2014; Roux et al. 2014; Seehausen et al. 2014; Wolf and Ellegren 2017; Johri et al.). For instance, recent evidence indicates that phylogenetic signal (e.g., the support for a particular topology) is associated with chromosome size and recombination rate, with larger chromosomes having slower rates and higher support for the species trees (Martin et al. 2019). However, most methods used in phylogenomics do not account for the multiple processes that shape the genomic landscape, which may confound estimation of evolutionary histories (Ewing and Jensen 2016; Schrider et al. 2016; Li et al. 2019). This is critical given that modern biogeography relies heavily on phylogenetic and population genetic approaches to explore the spatial history and demography of populations (Knowles 2009). Understanding how genomic architecture may affect inferences of spatial diversification histories will provide a clearer picture on the relative roles of intrinsic genomic characteristics, natural selection, and neutral processes on speciation (Pouyet et al. 2018; Johri et al. 2020).

Linked selection can have a large impact on genome-wide variation, but its effects on phylogenetic signal and demographic history of species are only starting to be explored (Martin et al. 2019; Li et al. 2019). The indirect influence of selection on linked neutral sites can reduce genetic diversity around target regions, decreasing local effective population size (*Ne*), and leading to faster fixation of alleles (Charlesworth 1998; Cruickshank and Hahn 2014; Burri et al. 2015). The intensity of linked selection on neutral sites is predicted by the interplay between the local density of targets under selection and the recombination rate, with more pronounced reductions in genetic diversity occurring in genomic regions with stronger selection and lower recombination (Smith and Haigh 1974; Hudson and Kaplan 1995; Zeng 2013; Gillespie 2000; Charlesworth et al. 1993). Areas of low recombination should also be more resistant to the confounding effects of gene flow and function as hotspots of phylogenetic signal (Chase et al. 2021; Martin et al. 2019). In these regions, linkage is maintained between introgressed variants and large genomic blocks may be removed from a population if deleterious alleles are present (Mořkovský et al. 2018; Brandvain et al. 2014; Schumer et al. 2017). The reduced impact of gene flow in regions of low recombination indicates that the phylogenetic signal is more likely to follow a bifurcating tree model, fitting the assumptions of most phylogenetic methods (Martin et al. 2019; Li et al. 2019). However, linked selection on low recombination areas violates neutral models of evolution and affects genome-wide estimations of demographic parameters (Schrider et al. 2016; Johri et al. 2020).

Although recent studies show that linked selection impacts a larger proportion of the genome than previously thought (Pouyet et al. 2018; Kern and Hahn 2018), the degree of this impact varies between species (Tigano et al. 2021; Jensen et al. 2019). For instance, the divergence between populations with high rates of gene flow might be restricted to small areas of the genome, maintained by strong divergent selection whereas the vast majority of the genome might show reduced differentiation due to widespread introgression (Ellegren et al. 2012). In contrast, genomic differentiation between populations with reduced levels of gene flow tends to be more widespread, given the higher contribution of genetic drift in isolated populations. This latter scenario should produce a stronger association between genomic architecture and levels of genetic differentiation across the genome.

In this study, we model the impact of genomic architecture on patterns of genetic diversity and spatial differentiation of three bird species that co-occur in southeastern Amazonian. These taxa have different propensities to move across space that are linked to their life histories, resulting in landscapes of genomic differentiation impacted by distinct levels of gene flow. We hypothesize that if linked selection led to congruent patterns of genetic diversity across the genome, then metrics associated with species differentiation and genetic diversity should be correlated with recombination rate, the density of targets under selection, and chromosome size. Alternatively, species could have idiosyncratic patterns of association with genomic architecture, driven by other factors such as historical demography and the level of differentiation across rivers. We demonstrate that the interplay between recombination, selection, and gene flow lead to a highly variable landscape of genetic diversity and differentiation within and between species, and impact evolutionary inferences under different population histories.

## Results

### Population genetics summary statistics and genomic features vary between species and across the genome

We resequenced 95 whole-genomes for three species of birds, *Phlegopsis nigromaculata* (n = 31), *Xiphorhynchus spixii* (n = 31), and *Lipaugus vociferans* (n = 26) that are co-distributed across three Amazonian areas of endemism, the Tapajos, Xingu, and Belem (Figure 1; Table S1). We obtained a mean coverage of 10.1x across all species. On average, 88% of the pseudo-chromosome reference genomes were recovered with coverage above 5x per individual (Table S1). Benchmarking Universal Single-Copy Orthologs analyses performed in BUSCO v2.0.1 (Waterhouse et al. 2018) identified a high proportion of targeted genes on the references used for *P. nigromaculata* (89.3%)*, X. spixii* (89.1%), and *L. vociferans* (93.4%; Table S2). The number of segregating sites were of a similar magnitude but varied between species: *P. nigromaculata* (n = 20,838,931), *X. spixii* (n = 26,583,784), and *L. vociferans* (n = 21,769,167). The proportion of missing sites per individual was on average 18% (Table S1). Summary statistics estimated from 100kb non-overlapping sliding windows and mean values per chromosome showed that levels of genetic diversity varied substantially across species and within and between chromosomes (Figure 2). Populations from species with higher putative dispersal abilities (*L. vociferans* and *X. spixii*) had greater nucleotide diversity (Figure 3; Tables S3-S5). We observed higher nucleotide divers!ity on smaller chromosomes in *P. nigromaculata* (Pearson’s correlation R = -0.6; p-value = 0.002; n = 26) and *X. spixii* (Pearson’s correlation R = -0.36; p-value = 0.047; n = 32) but not in *L. vociferans* (Pearson’s correlation R = -0.01; p-value = 0.94 ; n = 32; Figure S1-S6; Table S6-S11). We also found similar associations with *Dxy*, number of segregation sites, and Tajima’s D (Figure S1-S6; Table S6-S11). These results support a highly heterogeneous landscape of genetic diversity across the genome of the three studied species.

**Figure 1:**
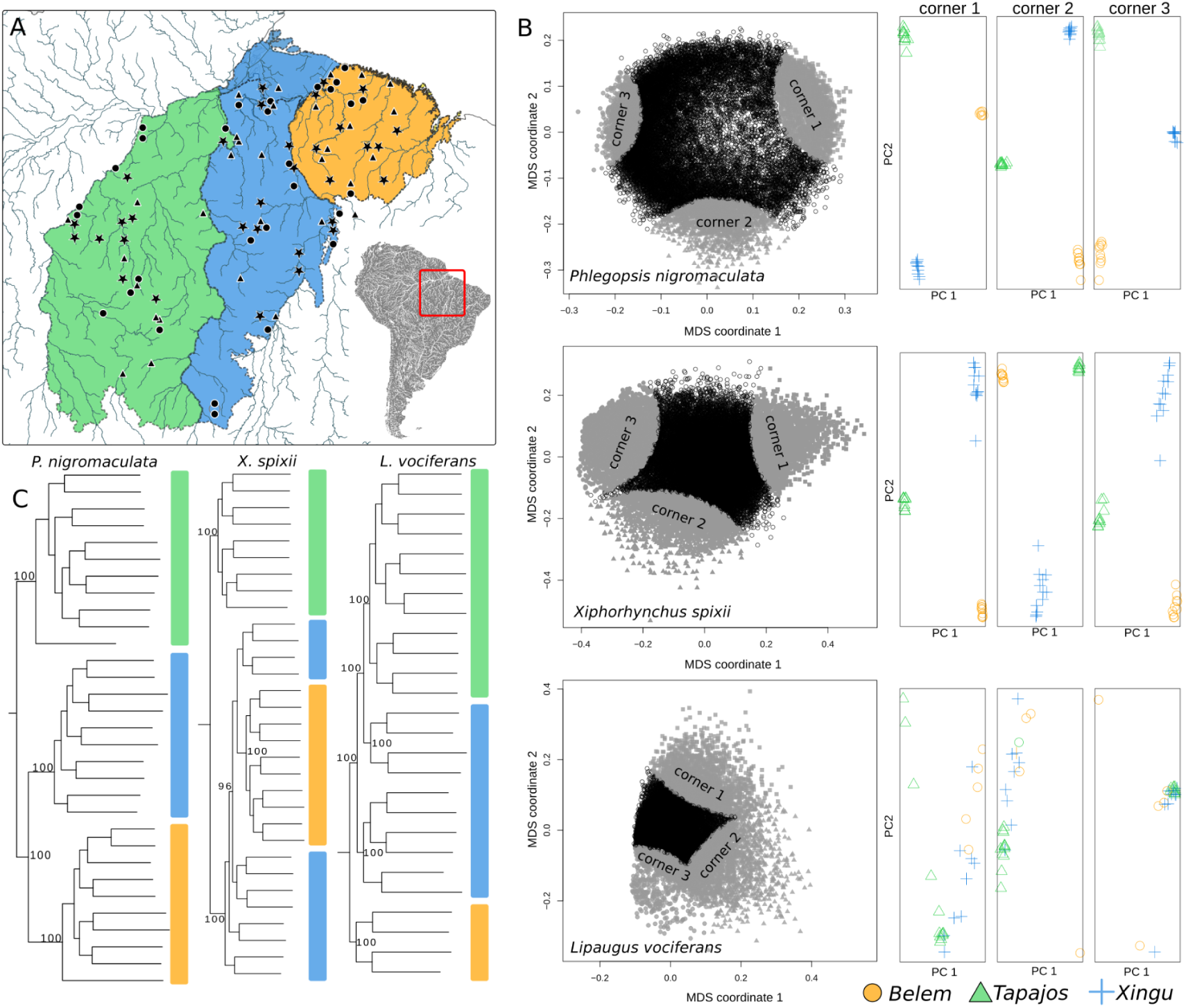
Contrasting patterns of genomic differentiation and spatial relationships between populations of three species of birds occurring in southeastern Amazonia. Geographic distribution of genomic samples for each species **(A)**. Triangles, stars, and circles are sampled localities for *Phlegopsis nigromaculata, Xiphorhynchus spixii,* and *Lipaugus vociferans,* respectively. Each colored polygon in the map represents a major Amazonian interfluve (area of endemism): Tapajos (Green), Xingu (Blue), and Belem (Yellow). **(B)** Patterns of genetic structure across the genome were obtained with local PCAs based on 10kb windows. Left: plots for the first and second multidimensional coordinates, where each point represents a genomic window. Gray points represent corners clustering the 10% of the windows closer to the three further points in the graph. Right: PCA plots for the first and second principal components, combining the windows of each corner. **(C)** Supermatrix phylogenetic estimations based on concatenated SNPs. Numbers on the nodes represent bootstrap support for major nodes in the tree. Color bars next to terminals represent geographic location following the map (A).

**Figure 2:**
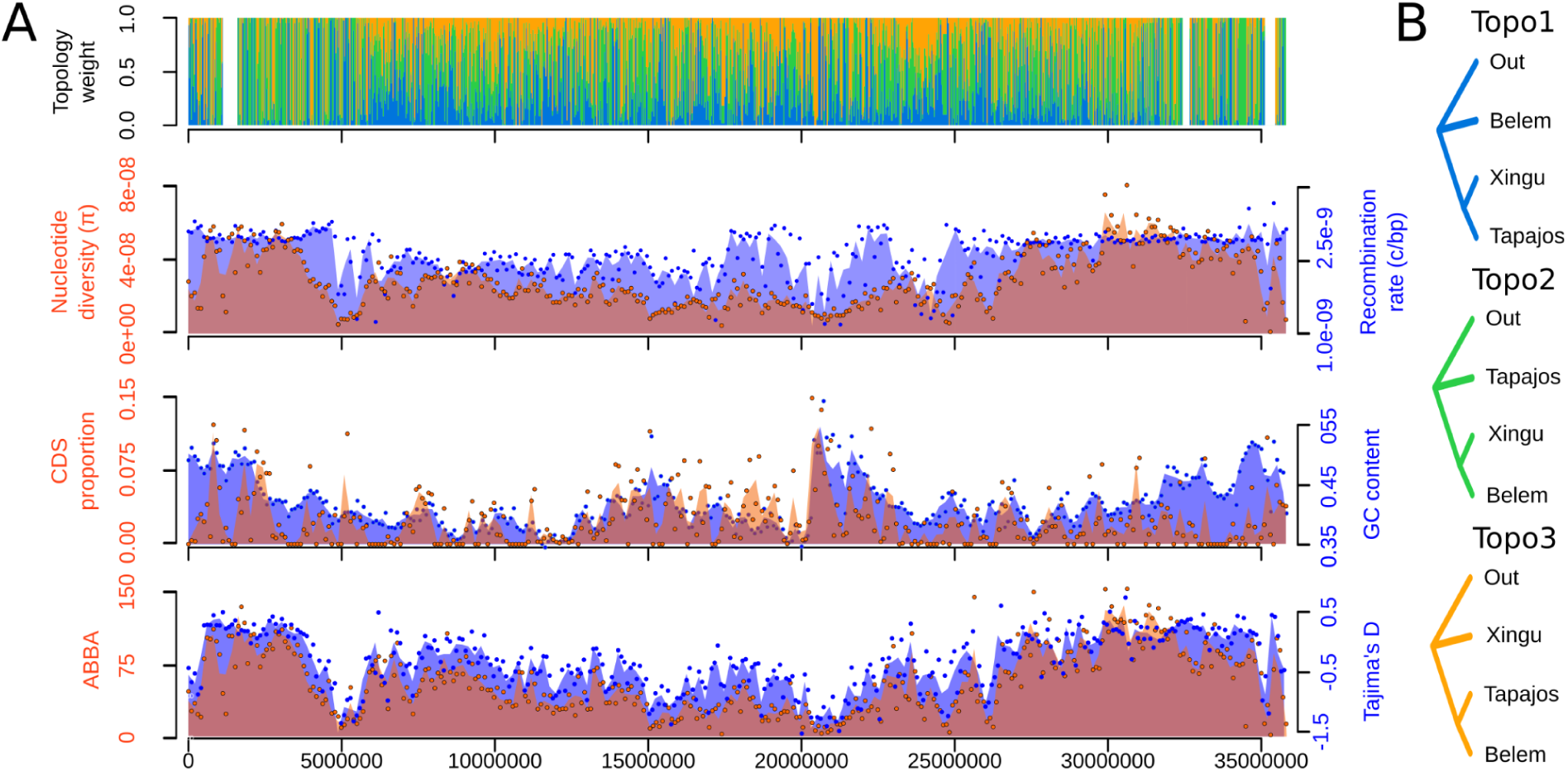
Phylogenetic signal for the species tree was higher on central portions of chromosomes and was associated with genomic architecture. **(A)** Example of how phylogenetic signal and summary statistics are distributed across a chromosome. Shown are pseudo-chromosome 6 of *P. nigromaculata*. On the top graph, colored bars represent the weight for the three alternative topologies shown in **(B)** for the relationship between Tapajos, Xingu, and Belem areas of endemism. On the three bottom graphs, the magenta color represents the overlap between the orange (y-axis on the left) and blue (y-axis on the right) tones. Estimates of nucleotide diversity, recombination rate, and Tajima’s D were based on the Tapajos population. ABBA represents the number of sites supporting Topology 2 assuming Topology 1 as the species tree.

**Figure 3:**
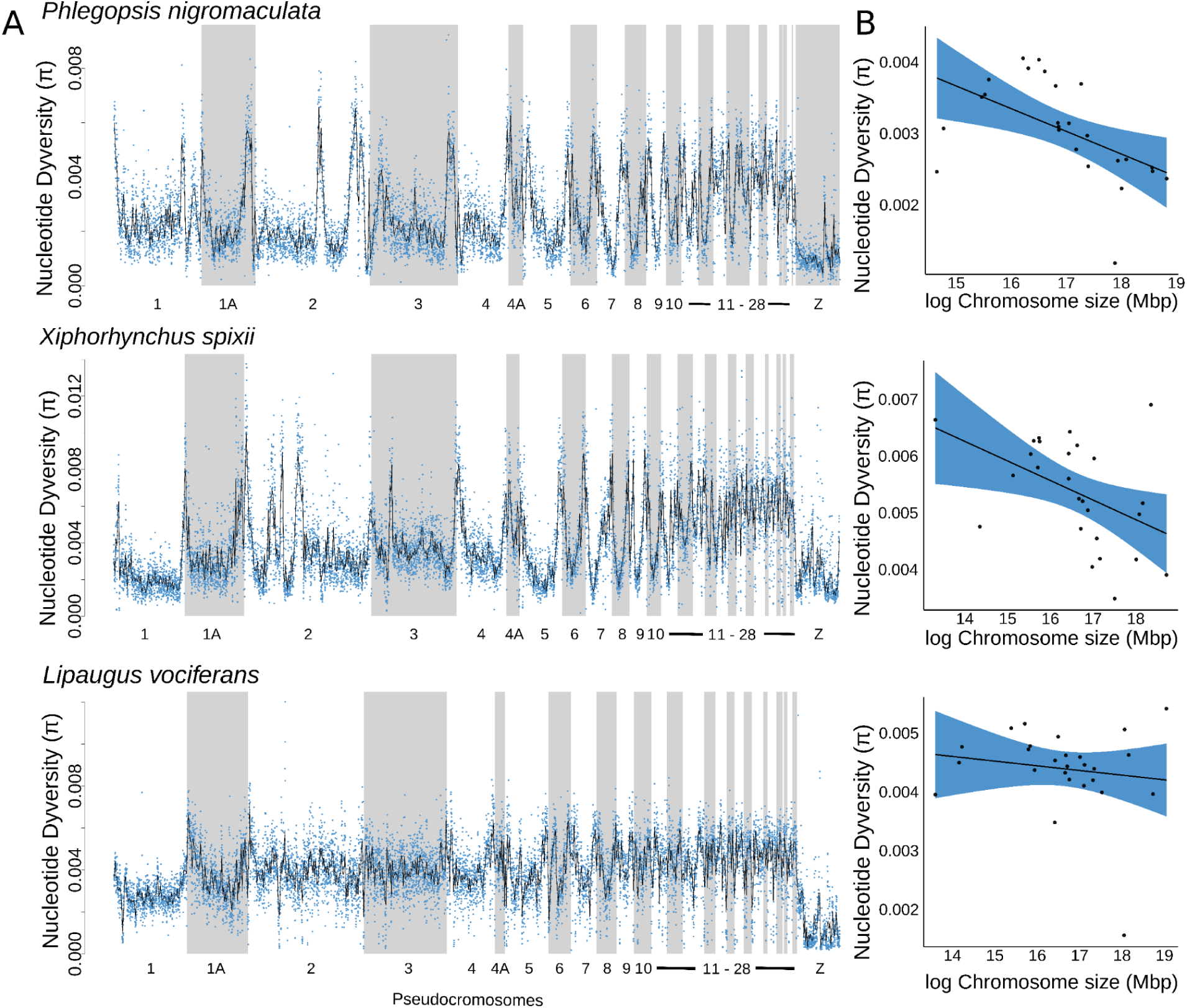
Nucleotide diversity varied within and between pseudo-chromosomes and across species. **(A)** Distribution of nucleotide diversity (π) across chromosomes for the three studied species.) across chromosomes for the three studied species. **(B)** Scatterplot and regression line with 95% confidence interval models with average nucleotide diversity as a function of chromosome size. *Phlegopsis nigromaculata*: Pearson’s correlation R = -0.6; p-value = 0.002; n = 26; *Xiphorhynchus spixii*: Pearson’s correlation R = -0.36; p-value = 0.047; n = 32; *Lipaugus vociferans*: Pearson’s correlation R = -0.01; p-value=0.94 ; n = 32.

We found that genomic regions with a reduced meiotic recombination rate were less impacted by gene flow and had stronger signatures of linked selection with greater genetic differentiation. To test for associations between recombination rate and genetic metrics while accounting for historical demography, we estimated the per-base recombination rate (r) with ReLERNN (Adrion et al. 2020). Recombination rate varied considerably across the genome of *P. nigromaculata* (mean r = 2.103e-9; SD = 4.413e-10), *X. spixii* (mean r = 1.234e-9; SD = 7.190e-10), and *L. vociferans* (mean r = 1.776e-9; SD = 4.025e-10; Figure S7) but in predictable ways. We found that regions with higher recombination rates were often in chromosome ends (Figure S7) and smaller chromosomes (Figure S4-S6) and were positively correlated with gene density and nucleotide diversity in all three species (Figure S8; Table S12). Loess models with recombination rate and gene density as covariate predictors explained a large proportion of the variation in genetic diversity in *P. nigromaculata* (R^2^ = 0.33), *X. spixii* (R^2^ = 0.65), and *L. vociferans* (R^2^ = 0.41; Figure S8, S9). These results suggest a significant effect of linked selection driving genomic patterns of diversity.

Genome-wide levels of differentiation between species match the evolutionary expectations associated with their life history. The least dispersive species that inhabit the understory, *P. nigromaculata*, had the most pronounced levels of genetic structure across rivers, followed by *X. spixii* which occupies the midstory, and the most dispersive, canopy species, *L. vociferans*, had the shallowest structure. To visualize patterns of genetic structure based on independently evolving sets of SNPs (linkage disequilibrium R^2^ < 0.2), we used Principal Component Analysis (PCA). Three isolated clusters of individuals supported strong geographic structure within *P. nigromaculata*, consistent with previous studies based on mtDNA, spatially matching areas of endemism (Silva et al. 2019; Aleixo et al. 2009b); Figure 1, S10). In *X. spixii*, the PCA supported strong differentiation between the Tapajos from Belem and Xingu populations, which had a substantial overlap in PC2 (5.0% of the explained variance; Figure S10). For *L. vociferans*, all samples clustered together, indicating a lack of spatial structure in the genetic variation (Figure S10). In agreement with these results, the average *Fst* between populations was considerably higher in *P. nigromaculata* (mean *Fst* = 0.1262; SD = 0.09) than in *X. spixii* (mean *Fst* = 0.059; SD = 0.046) and *L. vociferans* (mean *Fst* = 0.008; SD = 0.019). Pairwise *Fst* between populations was, in general, negatively correlated with genetic diversity metrics in all three species, and it was negatively correlated with recombination rate in *P. nigromaculata* and *X. spixii* (Figure S1–S6; Table S6–S11).

Genetic structure varied substantially across the genome and was associated with intrinsic genomic processes. To explore the genome-wide variation in genetic structure, we used local PCAs across sliding windows using lostruct v0.0.0.9 (Li and Ralph 2019). Local PCAs showed that distinct parts of the genome support different clustering patterns in *P. nigromaculata* and *X. spixii* (Figure 1). In *L. vociferans*, we observed a gradient between Tapajos, Xingu, and Belem individuals, without clear geographic structuring, consistent with the low *Fst* estimates reported for this species (Figure 1). Genetic structure, as described by the first two MDS axes obtained from lostruct, were associated with recombination rate in *P. nigromaculata* (MDS1: R^2^ = -0.04, p-value < 0.0001, n = 20,143 windows) and *X spixii* (MDS1: R^2^ = 0.017, p-value < 0.0001, n = 28,803 windows; MDS2: R^2^ = -0.15, p-value < 0.0001, n = 28,803 windows) but not in *L. vociferans* (R^2^ < 0.001 for all MDSs, n = 25,007 windows). This result indicates that for the species with marked genetic structure across rivers, recombination rate was a key predictor of spatial differentiation. These results highlight the high variation in patterns of genetic structure across the genome as well as the contrast between patterns of diversification of sympatric species distributed across Amazonian rivers.

Although the association between recombination rate and genetic diversity supports the effect of linked selection, it does not indicate which portions of the genome are directly impacted by this process. To further explore the extent of linked selection across the genome, we used a machine learning approach implemented on diploS/HIC (Kern and Schrider 2018) to predict which 20kb genomic windows were evolving under neutrality or had signatures of selective sweeps and linked selection (i.e., background selection; Charlesworth et al. 1993). We initially simulated genomic windows under distinct selective regimes accounting for historical oscillations in effective population size and uncertainty in demographic parameters using discoal (Kern and Schrider 2016). To account for the historical demography of the analyzed populations in our simulations, we calculated population size changes occurring in the last 300,000 years with SMC++ (Terhorst et al. 2017); Figure S11). The convolutional neural network designed to classify five alternative models produced an average accuracy for model classification of 0.69 and a false positive rate of 0.27 among species (Table S13). On average, across all three species, 43.3% of the genome had signatures of selective sweeps or linked selection. In *P. nigromaculata*, 30.29% of tested windows had a high probability (>0.70) for models including the direct or indirect effect of selection (Figure 4). For *X. spixii* (44.83%) and *L. vociferans* (54.77%), we obtained even higher proportions (Figure 4). When accounting for a false positive rate by assuming that all potential false positives were neutral regions classified as having signatures of selection, on average 31.6% of the genomes we analyzed had signatures of selective sweeps or linked selection.

**Figure 4:**
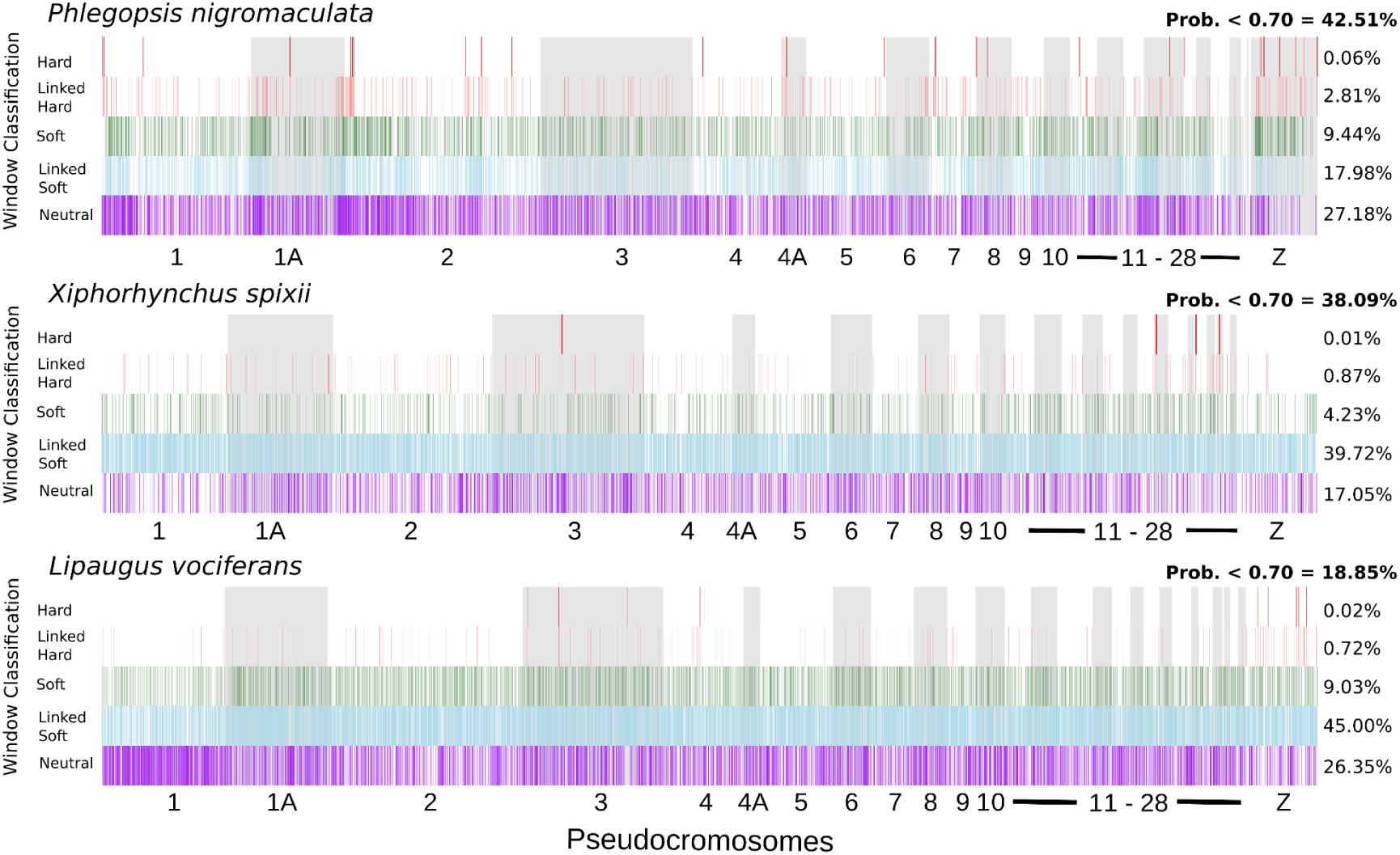
Signature of selection across the genomes of the studied species. Vertical bars represent the model with the highest probability for 20kb genomic windows. On the right is the percentage of windows assigned to each of the five models with high probability (>0.70): hard sweep, linked to hard sweep, soft sweep, linked to soft sweep, and neutral. In Bold is the proportion of windows with low probability for model classification.

### Phylogenetic signal was associated with genomic architecture

We explored how evolutionary relationships were distributed across the genome of the co-occurring species to test which aspects of the genomic architecture best predicted phylogenetic signal. First, we estimated topologies for each species in IQTREE-2 v2.1.5 (Nguyen et al. 2015) by concatenating SNPs and controlling for ascertainment bias (Figure 1). We found substantial variation in topology between species, with clades matching the three areas of endemism only in *P. nigromaculata.* In *X. spixii*, Belem individuals were nested within Xingu despite forming a monophyletic group. In *L. vociferans*, the clustering of individuals matched their spatial distribution.

The support for alternative species tree topologies varied with recombination rate and genetic diversity (Figure 5). To explore how phylogenetic relationships varied with genomic characteristics and population genetics summary statistics, we calculated gene trees for non-overlapping genomic windows and ran species tree analyses for subsets of the genome. For *P. nigromaculata* we did not obtain high support for any topology in the genome-wide species tree analysis, but the topology with the highest probability (posterior probability = 0.81) matched the concatenated SNP tree (Figure 5). The topology estimated from genomic regions with high recombination matched the concatenated tree, but regions of low recombination placed Tapajos and Xingu as sisters (Figure 5). A similar pattern was observed when filtering gene trees based on π) across chromosomes for the three studied species. and *Dxy*. The phylogenetic signal in *X. spixii* and *L. vociferans* were more stable, with widespread support for the same topology across the genome but with substantially higher weight for that topology in areas with lower recombination and lower genetic diversity. Phylogenetic signal also co-varied with chromosome size in *P. nigromaculata* but not in *X. spixii* and *L. vociferans*. In *P. nigromaculata*, macro chromosomes supported the topology found in low recombination areas (Topology 1), while microchromosomes (<50MB) supported the concatenated tree (Topology 2; S12- S14). Our results suggest that support for the species tree was higher in genomic regions with reduced genetic diversity and recombination rates.

**Figure 5:**
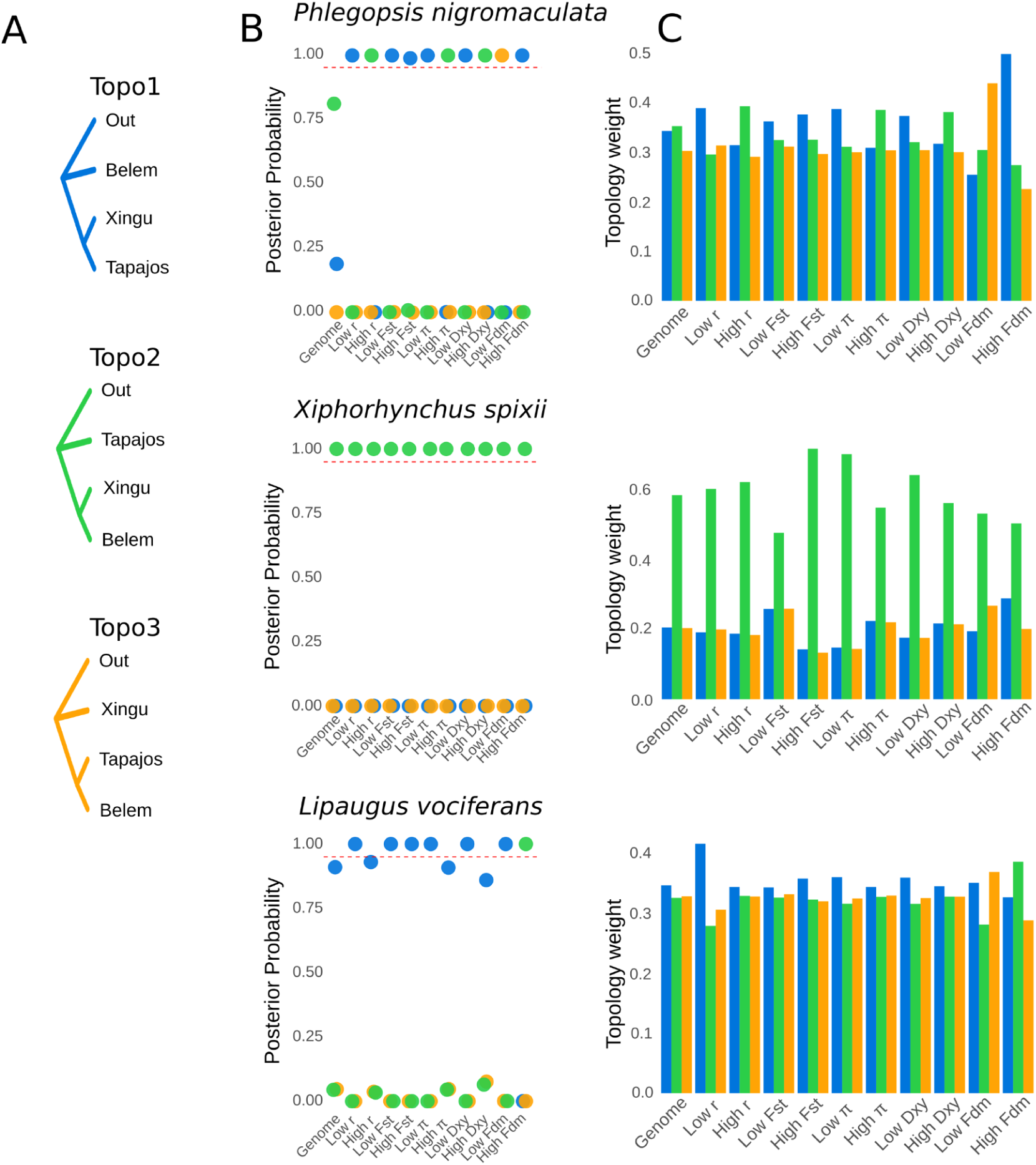
Species tree and topology weights vary accordingly to recombination rate and genetic diversity. **(A)** Alternative topologies for the relationship between the three areas of endemism and the outgroup; **(B)** posterior probabilities for the three topologies for windows across the whole-genome and for distinct subsets of genomic windows that were selected based on upper and lower thresholds for summary statistics; and **(C)** weights for three topologies for windows across the whole-genome and for distinct subsets of genomic windows that were selected based on upper and lower thresholds for summary statistics: - recombination rate (*r)*; fixation index (*Fst*); nucleotide diversity (*π*); genetic distance (*Dxy*); and introgression proportion (*Fdm*).

The weight for alternative topologies varied considerably across windows and was associated with genomic architecture. To test how the probability of alternative topologies varied across the genome of each species, we calculated topology weights using Twisst (Martin and Van Belleghem 2017). This analysis was performed on genomic windows with 100 SNPs and assumed the three possible unrooted trees representing the relationship between the three areas of endemism plus an outgroup. From hereafter we refer to these three unrooted topologies as Topology 1 (outgroup, Belem (Xingu, Tapajos)), Topology 2 (outgroup, Tapajos (Xingu, Belem)), and Topology 3 (outgroup, Xingu (Belem, Tapajos)). When averaging weights for genome-wide windows of *P. nigromaculata* we observed a higher weight for Topology 2, followed closely by Topology 1. Genomic windows from *P. nigromaculata* based on upper and lower thresholds for summary statistics showed substantial variation in which topology had the highest average weight, consistent with our species tree approach (Figure 5). For the other two species, we found less variation along the genome for the topology with the highest average weight. The topology with the highest weight also varied across chromosomes of different sizes. Smaller chromosomes in *P. nigromaculata* had a higher weight for Topology 2, putatively derived from gene flow (Figure 6). In *X. spixii* we observed a progressive increase in the weight for the species tree (Topology 2) in larger chromosomes, despite a non-significant correlation, whereas in *L. vociferans* all three topologies had a similar weight across chromosomes.

**Figure 6:**
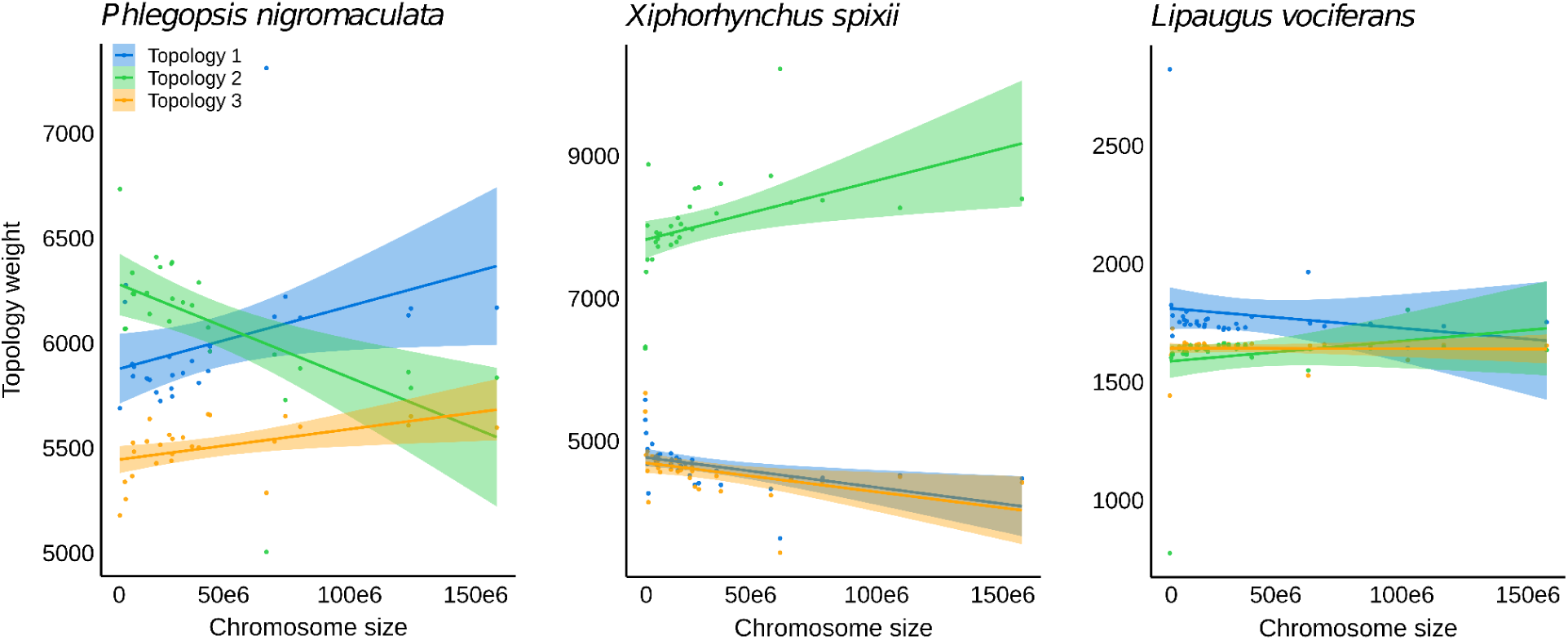
Chromosome size was associated with topology weight across the genome of *Phlegopsis nigromaculata* (Topology 2 - R^2^ = 0.31, p-value = 0.003; Topologies 1 and 2 p-value > 0.05, n= 26) but not in *Xiphorhynchus spixii* (p-value > 0.05 for all three topologies, n = 32), and *L. vociferans* (p-value > 0.05 for all three topologies, n = 32). Scatterplot and regression line with 95% confidence interval showing the relationship between topology weights and chromosome size. We tested three alternative unrooted topologies for the relationship between the three areas of endemism (Tapajos, Xingu, and Belem): Topology 1 (outgroup, Belem (Xingu, Tapajos)), Topology 2 (outgroup, Tapajos (Xingu, Belem)), and Topology 3 (outgroup, Xingu (Belem, Tapajos)).

The conflicting phylogenetic pattern observed for *P. nigromaculata* could be driven by gene flow increasing the signal for the topology where introgressing populations are sisters. To explore how topology weight varied according to gene flow and intralocus recombination, we performed coalescent simulations with demographic parameters similar to those estimated for *P. nigromaculata,* and we calculated topology weights using the approach mentioned above. Our simulations suggested that in the absence of gene flow, the frequency of alternative topologies was similar (Figure S15). The presence of gene flow between non-sister species produces a deviation from this pattern by increasing the average weight for the topology with introgressing populations as sisters. This relationship was further intensified by intralocus recombination (Figure S15). Although our simulations corroborate that recombination rate by itself does not affect levels of genetic diversity (Hudson 1983), it does affect levels of ILS between populations when gene flow was present by increasing the variance of topology weights across the genome (Figure S15). When comparing the results obtained with this simulation approach with the genome-wide topology weights obtained for *P. nigromaculata,* our results suggest that gene flow between non-sister taxa was likely increasing the weights for one of the two best topologies.

### Gene flow affected phylogenetic inference

When modeling gene flow, our results indicated that the topology recovered for low recombination areas was the best genome-wide tree. To estimate the probability for alternative topologies and demographic parameters for the entire genome explicitly accounting for gene flow, we used a multiclass neural network approach with Keras v2.3 (https://github.com/rstudio/keras) in R. We simulated genetic data under the three possible unrooted topologies for the relationship between areas of endemism using uniform priors for *Ne*, gene flow between geographically adjacent populations, and divergence times. We selected one 10kb window every 100kb to reduce the effect of linkage between windows, excluding windows with missing data. This procedure yielded a total of 7,213, 9,140, and 9,693 windows for *P. nigromaculata, X. spixii,* and *L. vociferans,* respectively, and we randomly selected 5,000 windows per species. Genomic windows were converted into feature vectors representing the mean and variance of commonly used population genetics summary statistics. On average, this approach produced highly accurate model classification probabilities (neural network accuracy = 0.93; categorical cross-entropy = 0.17) and a high correlation between observed and estimated parameters for testing data sets with low mean absolute errors (Table S14-S16). PCAs and goodness-of-fit analyses showed that simulated models matched observed values of summary statistics. In *P. nigromaculata,* we obtained a high probability for Topology 1 (probability = 0.86), conflicting with the concatenated and species tree topology (Topology 2; probability = 0.12) but agreeing with the topology of low recombination areas (Table S17). Divergence times were highly variable between species. The initial divergence in *P. nigromaculata* between Belem and the ancestor of the Tapajos and Xingu lineages, diverged at 149,836ya (SD = 15,272; MAE = 36,576; Table S14), followed by the divergence between the later populations at 77,866ya (SD = 17,849; MAE = 37,810). In *X. spixii* (Topology 2; probability = 0.99), the first divergence event occured at 218,858ya (SD = 12,095; MAE = 30,668), followed by a more recent divergence event at 40,303ya (SD = 16,236; MAE = 32,798; Table S15). For *L. vociferans* (Topology 1; probability = 0.54) divergence times were the most recent, occurring within the last 40,000 years, reflecting the lack of population structure in this species (Table S16). Our data indicated that gene flow among *P. nigromaculata* populations (2Nm) was negligible between Tapajos and Xingu (migration between Tapajos and Xingu = 0.002; SD = 0.005; MAE = 0.161) and low between the non-sisters in Xingu and Belem (migration between Xingu and Belem = 0.484; SD = 0.486; MAE = 0.138; Table S14). The relatively reduced levels of gene flow between populations of *P. nigromaculata*, indicated that ancestral gene flow might be the source of the phylogenetic conflict. In *X. spixii*, we inferred moderate rates of gene flow between populations, which was highest between the recently diverged Xingu and Belem populations (migration between Xingu and Belem = 2.075; SD = 0.144; MAE = 0.139; Table S15). In *L. vociferans* we also found moderate to high gene flow among populations (migration between Tapajos and Xingu = 2.347; MAE = 0.175; migration between Xingu and Belem = 1.827; MAE = 0.205; Table S16).

### Selection biases estimates of demographic parameters

Given the considerable proportion of the genome with signatures of selective sweeps and background selection in all three species, we explored how selection might impact demographic parameters. We calculated parameters from subsets of genomic windows classified under distinct selection regimes with diploS/HIC using our machine learning approach. We selected up to 1,000 windows assigned to each of the five models tested in diploS/HIC with a probability > 0.70, and estimated demographic parameters based on the topology with the highest probability considering all genomic windows (Topology 1 for *P. nigromaculata* and *L. vociferans*, and Topology 2 for *X. spixii*). We found that genome-wide windows yielded more similar values for *Ne* and gene flow to regions with signatures of linked selection than neutral regions (Figure 7). Our approach supported higher *Ne* and gene flow in neutral areas of the genome than areas under selection with little to no overlap of standard error distributions (Figure 7).

**Figure 7:**
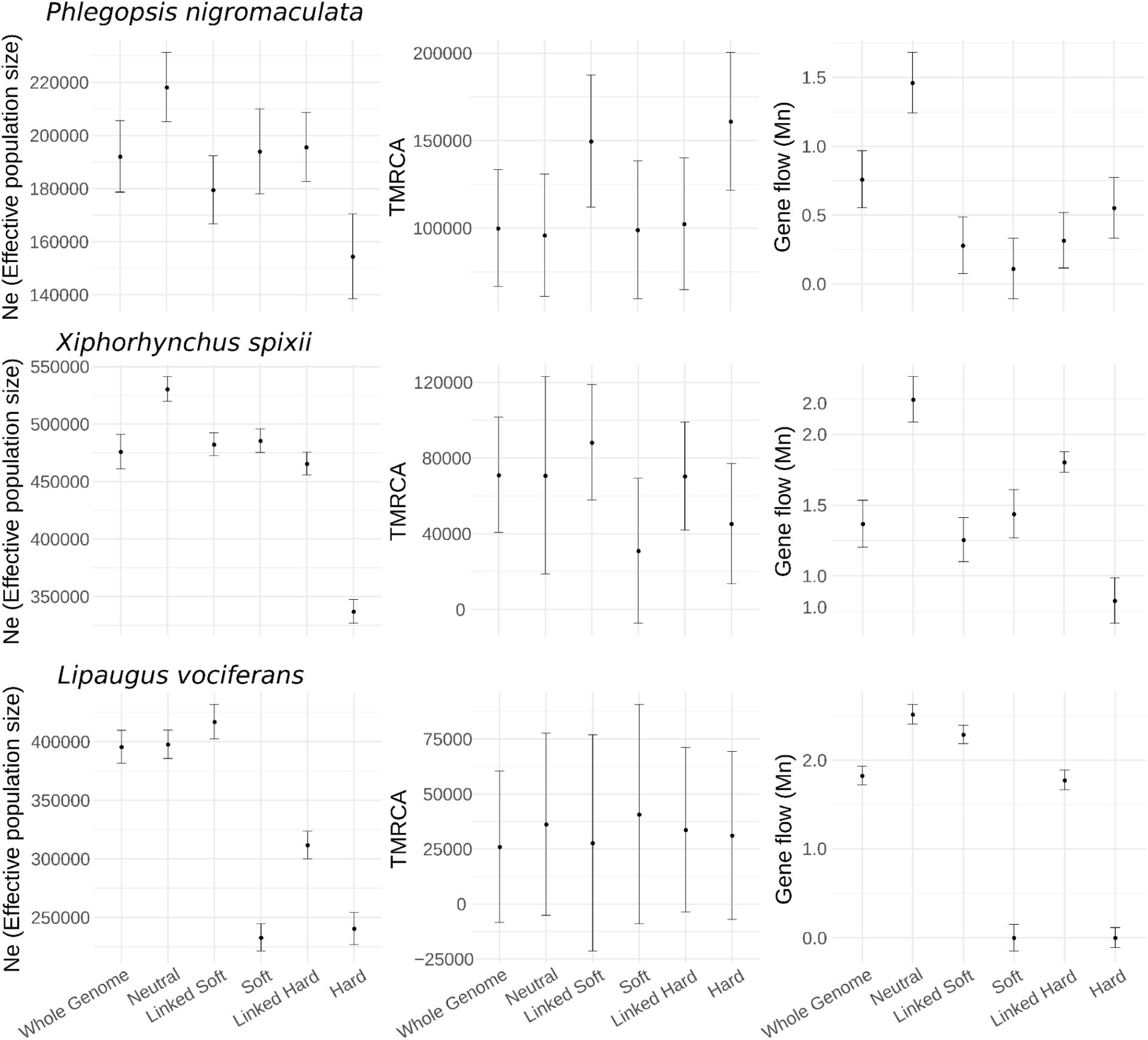
Genomic regions inferred to be evolving neutrally had larger effective population sizes and higher gene flow than areas with signatures of selection. Demographic parameter estimation for the three studied species of Amazonian Birds. - Effective population size of the Tapajos population (*Ne*). - Time to the most recent common ancestor of the most recent divergence event (TMRCA). - Gene flow rate between Tapajos and Xingu populations (Gene flow). Classes on the x-axis represent genome-wide windows (Whole Genome), and subsets of genomic windows assigned with high probability to distinct models tested with diploS/HIC. Neutral: neutrally evolving windows; Linked Soft: windows linked to a soft sweep; Soft: windows assigned to a soft sweep; Linked Hard : windows linked to a hard sweep; and Hard: windows assigned to a hard sweep.

Demographic parameters varied considerably across the genome and were strongly associated with the recombination rate (Figure S16; Table S18). To explore the associations between demographic parameters and genomic architecture, we calculated the probability of alternative topologies and estimated demographic parameters for 100kb genomic windows, taking into account intralocus recombination. To increase model classification accuracy, we only tested the two most likely topologies based on the spatial distribution of the populations. This approach yielded a high accuracy in model classification (accuracy = 0.9314; categorical cross-entropy = 0.18). We also recovered high correlations between simulated and pseudo observed data indicating good accuracy in parameter estimation for *Ne* (average R^2^ = 0.94; average MAE = 73,099 individuals) and divergence times (average R^2^ = 0.87; average MAE = 39,172ya) but not for gene flow (average R^2^ = 0.54; average MAE = 0.19 migrants per generation). Effective population sizes and divergence times varied over one order of magnitude and gene flow over two orders of magnitude across the genome. The substantial variation in *Ne* and gene flow across the genome was positively associated with recombination rate in all three species, except gene flow in *L. vociferans* (Figure S16; Table S18). Variation in divergence time was not associated with recombination rate in any of the species (Table S18).

To further explore the signal for gene flow across the genome, we estimated D and fdm statistics using window-based ABBA-BABA tests in 100kb non-overlapping sliding windows. We found little evidence for gene flow between populations of all three species, except between Belem and Xingu populations of *P. nigromaculata.* The gene flow inferred between these two populations of *P. nigromaculata* was positively associated with recombination rate (Figure 2, S1, S4). For *L. vociferans* despite the lack of genetic structure, the D statistics failed to find any significant levels of introgression, likely due to the high levels of gene flow among all three populations, violating the ABBA-BABA model.

## Discussion

We found that genomic architecture directly impacts genome-wide estimates of key parameters used in speciation research, adding an underappreciated layer of complexity for testing diversification hypotheses. Biogeographic patterns inferred from Amazonian taxa exhibit a wide array of temporal and spatial divergences (Silva et al. 2019; Smith et al. 2014; Dagosta and De Pinna 2019; Lynch Alfaro et al. 2015; Byrne et al. 2018; de Oliveira et al. 2016; Penz et al. 2015). Our results indicate that the heterogeneity in parameters estimated in these studies may be driven by unaccounted for evolutionary processes. We found that the interplay of selection, gene flow, and recombination shaped the genomic landscape of genetic diversity resulting in different portions of the genome strongly supporting alternative hypotheses on spatial diversification. This study indicates that genome-wide estimates of phylogenomic parameters might be misleading and accounting for the processes that produce an heterogeneous genomic landscape is essential to understand the processes driving speciation across Amazonian rivers.

### Genomic architecture informs patterns of spatial diversification

We found that introgression, which was associated with genomic architecture, can produce a highly heterogeneous landscape of phylogenetic conflict that obscures inferences on population history. For example, in *P. nigromaculata*, gene flow between non-sister taxa was positively associated with recombination rate and the support for an alternative topology (topology 2) that was found to be the most prevalent tree across the genome. This pattern affected genome-wide species tree and topology weight analyses. In contrast, the topology reflecting the most probable species tree (topology 1) was more frequent in windows with low recombination rates. The phylogenetic conflict between alternative topologies has practical implications for inferring the pattern of spatial population history. Support for topology 2 for *P. nigromaculata* would indicate that the taxa diverged via a stepping-stone process from the west through the Tapajos, Xingu, and Belem regions, consistent with the Moisture Gradient Hypothesis (Silva et al. 2019). In contrast, if topology 1 reflects the population history of *P. nigromaculata*, it would indicate an opposite scenario, with an ancestral population in southeastern Amazonia, which could be linked to physiographic changes in the landscape (Musher et al. 2021; Albert et al. 2018). Although the probability for alternative topologies was more stable across the genomes of *X. spixii* and *L. vociferans*, topology weight was also predicted by recombination rate and gene flow. The conflicting phylogenetic pattern reported here might be common across the thousands of lineages isolated by Amazonian tributaries, given that recent studies have been suggesting extensive introgression across rivers (Barrera-Guzmán et al. 2018; Berv et al. 2021; Ferreira et al. 2018; Del-Rio et al. 2021; Weir et al. 2015; Musher et al. 2021).

We found that the interaction between intrinsic (e.g. recombination and selection) and extrinsic (e.g., gene flow) genomic processes lead to phylogenetic conflict affecting genome-wide estimates. In the presence of gene flow, low recombination areas are more prone to maintain the ancient branching signal (Li et al. 2019; Tigano et al. 2021). The strong linkage in low recombination regions should lead to the more effective removal of alleles introduced by hybridization that are more likely to be deleterious (Nachman and Payseur 2012; Schumer et al. 2018). In this sense, even methods designed to incorporate gene flow into phylogenetic analyses such as phylogenetic network approaches (Wen et al. 2018; Solís-Lemus and Ané 2016), might produce misleading results when considering genome-wide markers. Although phylogenetic networks are an ideal way to track the presence of gene flow, it might be difficult to disentangle the processes driving phylogenetic conflict, given that introgression proportions might be biased by the genomic landscape.

### Selection and recombination effects parameter estimates

We showed pervasive signatures of selection across the genome of three co-occurring species impacting genome-wide demographic parameter estimation. On average and after accounting for false positives, one third of the genomes of our focal species had a high probability for models with selective sweeps or linked selection. Recent estimations for birds and mammals show substantial variation in the proportion of the genome subject to selection, with approximations above 50% (Manthey et al. 2021; Brand et al. 2021; Pouyet et al. 2018; McVicker et al. 2009). Although our data indicate that regions potentially affected by linked selection had a better fit to a bifurcating phylogenetic model, these regions violate neutral models of evolution, affecting genome-wide population genomic parameters (Schrider et al. 2016; Pouyet et al. 2018). Whole genome approximations of *Ne* and gene flow were more similar to regions with signatures of selection versus regions deemed to be evolving neutrally. For instance, areas of the genome evolving neutrally had up to 13% larger *Ne* in *P. nigromaculata* and 64% higher gene flow in *X. spixii* than estimates based on genome-wide loci (Figure 7). These results are in agreement with studies indicating that demographic parameters can be severely affected by positive and background selection (Schrider et al. 2016; Johri et al. 2021). Natural selection can skew levels of genetic variation in a similar way to certain non-equilibrium demographic histories, often leading to overestimates of population bottlenecks and the rate of demographic expansions (Schrider et al. 2016; Ewing and Jensen 2016). For example, positive selection leading to fixation of large haplotypes linked to the target of selection may mimic population bottlenecks (Wayne and Simonsen 1998), and the recovery from these sweeps, might inflate the proportion of rare variants, resembling recent population expansions (Schrider et al. 2016). Although we did not model demographic changes, summary statistics that are indicative of demographic oscillations such as Tajima’s D, varied considerably across the genome, with more negative values in regions of low recombination (Figure 2). Our examination of three codistributed species showed that the effect of selection on demographic analyses was a general phenomenon that can have a profound effect on modeling population histories.

Reconciling phylogenetic inference and demographic parameter estimation from whole genomes are aided by recombination and selection-aware approaches. By characterizing the genomic landscape, the effect of selection in demographic parameter estimations can be mitigated by targeting genomic regions as distant as possible from potential targets of selection such as genes and functional elements, as well as avoiding areas of low recombination or affected by biased gene conversion (Pouyet et al. 2018). A key problem with selecting loci with distinct characteristics is that current methods designed to calculate recombination and selection across the genome achieve optimal performance when the demographic history of a population is known (Harris et al. 2018; Dapper and Payseur 2018; Johri et al. 2020; Rousselle et al. 2018). On the other hand, demographic parameters may be heavily biased when recombination and selection are neglected (Pouyet et al. 2018; Ewing and Jensen 2016). This conundrum indicates that methods designed to simultaneously account for multiple genomic processes, such as recombination, selection, drift, and mutation (Barroso and Dutheil 2021; Johri et al. 2020, 2021), associated with simulation studies (Tigano et al. 2021) might be necessary to unbiasedly obtain evolutionary parameters from genome-wide variation.

### Heterogeneous genomic landscapes within and between species

The recombination rate and the proportion of the genome impacted by selection varied considerably among species leading to different levels of association between genomic architecture and population history. Chromosome size was a good predictor of genetic diversity, recombination rate, and phylogenetic signal in two of the species but not in *L. vociferans*. The lack of association between chromosome size and genomic characteristics in *L. vociferans* was unexpected given that during meiosis, chromosome segregation often requires at least one recombination event per homologous chromosome pair (Fledel-Alon et al. 2009). This process should lead to an overall higher recombination rate in shorter chromosomes (Manthey et al. 2021; Farré et al. 2012; Kawakami et al. 2014; Haenel et al. 2018; Kaback et al. 1992). Similar results to the one we obtained for *L. vociferans* have also been observed in other species of birds and mammals (Dutoit et al. 2017a; Pessia et al. 2012; Kartje et al. 2020) and could be explained by a reduced synteny between our reference and the zebra finch genome, or the historical demography of the species, which in some cases can reverse the expected associations between recombination, genetic diversity, and chromosome size (Tigano et al. 2021; Van Belleghem et al. 2018). However, the former scenario was less likely due to the relative stability of chromosomes across avian species (Ellegren 2010). These results suggest that, although genomic architecture seems to be a strong predictor for phylogenetic relationships in the presence of gene flow, this was not the case in *L. vociferans*.

Genomic architecture was likely conserved over the timeframe of our study, indicating that a stable genomic landscape might shape the demographic and phylogenetic histories of closely related populations in similar ways (Vijay et al. 2017; Van Doren et al. 2017; Dutoit et al. 2017b; Delmore et al. 2018; Tigano et al. 2021). For example, consistent variation in *Fst* values across the genomes of population pairs of the same species were likely reflecting the genomic landscape of the ancestral population. Regions of low recombination and *Ne* in the parent population would promote faster differentiation between daughter populations after isolation. These results agree with the idea that in birds, recombination hotspots are associated with gene promoters, which might help maintain a conserved landscape of recombination across lineages that span for millions of years (Singhal et al. 2015). Although the temporal scale on which the genomic architecture remains conserved between diverging lineages is still unknown, our data suggest that stable recombination hotspots in the early stages of speciation can considerably help accounting for genomic architecture in phylogenomic approaches.

### Multiple processes produce heterogenous landscapes across the genome

Identifying the driving forces shaping patterns of diversity along the genome of non-model organisms and understanding how theoretical models extend to natural systems is a major endeavor in speciation genomics (Comeron 2014; Elyashiv et al. 2016; Stankowski et al. 2019; Barroso and Dutheil 2021). In this study, we demonstrated that the interplay between recombination and selection had a strong impact on phylogenetic inference and demographic parameters, which are key for distinguishing alternative spatial models of divergence. However, other genomic processes might be shaping phylogenetic signals across the genome. For instance, levels of polymorphism across the genome could be derived from variation in mutation rate (Besenbacher et al. 2019; Jónsson et al. 2018; Smith et al. 2018; Barroso and Dutheil 2021). Non-crossover gene conversion (Korunes and Noor 2017), where DNA strands break during meiosis and are repaired based on homologous sequences without crossing-over, and crossover events could be mutagenic, leading to higher mutation rates in areas of higher recombination (Arbeithuber et al. 2015; Korunes and Noor 2017). Simulation studies have rejected gene conversion as a process driving genome-wide patterns of genomic diversity in relatively recent divergence events (Tigano et al. 2021) and empirical studies suggest that mutations associated with crossover events occur at relatively low frequencies (Halldorsson et al. 2019). Although differential mutation rates across the genome might not explain the strong association between genetic diversity and genomic architecture, the majority of the variation in genetic diversity in our focal species was not explained by recombination. This suggests that variation in mutation rate, not associated with recombination, could be playing a role in the genomic landscape of genetic diversity. It is important to note that irrespective of the processes driving the heterogeneous levels of genetic variation across the genome, these biases on genome-wide phylogenetic and population genetics inferences may remain unless the multitude of parameters varying across the genome are modeled in a unifying approach (Johri et al. 2021). Until that horizon is reached, genomic-architecture-aware approaches can be used to disentangle the effects of intrinsic genomic characteristics and selection from neutral processes.

## Methods

### Studied species, sampling design and whole-genome sequencing

We selected three species that occur in southeastern Amazonia occurring in distinct forest strata of upland forest habitats: 1) *Phlegopsis nigromaculata,* an obligatory army-ant follower restricted to the understory, with three distinct subspecies isolated by Xingu and Tocantins rivers with considerable levels of genetic differentiation (Aleixo et al. 2009a); 2) *Xiphorhynchus spixii,* which occupies the midstory of eastern Amazonian forests, and has two structured populations divided by the Xingu River (Aleixo 2004); and 3) *Lipaugus vociferans,* a widespread canopy species that is expected to be less structured across rivers.

To optimize the spatial representation of our samples, we selected a single individual per locality targeting approximately 10 individuals per interfluve per species (Tapajos, Xingu, and Belem), yielding a total of 31, 31, and 26 samples for *P. nigromaculata, L. vociferans*, and *X. spixii*, respectively (Table S1; Figure 1). We isolated genomic DNA from muscle tissue preserved in alcohol (n = 65) and skin from the toe pads of museum specimens (n = 31). All samples were loaned from the Museu Paraense Emilio Goeldi (MPEG). From tissues, we extracted DNA with Qiagen high molecular weight DNA kit (MagAttract HMW DNA Kit - Qiagen). For the toe pads, we performed a protocol specific for degraded DNA consisting of additional steps for washing the samples with H2O and EtOH prior to extracting and extra time for digestion. We modified the DNeasy extraction protocol (DNeasy Blood & Tissue Kits - Qiagen) by replacing the standard spin columns with the QIAquick PCR filter columns (QIAquick PCR Purification Kit - Qiagen), selecting for smaller fragments of DNA, typically found in degraded samples. Toe pad extractions were conducted on a dedicated lab for working with historical samples at the American Museum of Natural History (AMNH) to reduce contamination risk. We quantified DNA extracts using a Qubit 2.0 Fluorometer (Thermo Fisher Scientific). Illumina libraries with variable insert sizes were generated and samples were sequenced by Rapid Genomics (Gainesville, Florida) to ∼10x coverage using 3.5 lanes of paired-end (2x150 bp) Illumina S4 NovaSeq 6000. Raw reads were trimmed and filtered using trimmomatic v0.36 (Bolger et al. 2014).

### Genomic references, gene annotation and outgroups

We obtained reference genomes from closely related species. For *P. nigromaculata,* we used as reference the genome of *Rhegmatorhina melanosticta* (Coelho et al. 2019) with TMRCA = 9.60Ma (Harvey et al. 2020). For *X. spixii,* we used the genome of *X. elegans* (GCA_013401175.1 ASM1340117v1; NCBI genome ID: 92877; Feng et al. 2020) with TMRCA = 2.36Ma (Harvey et al. 2020), and for *L. vociferans* we used the genome of *Cephalopterus ornatus* (GCA_013396775.1 ASM1339677v1; NCBI genome ID: 92752; Feng et al. 2020) with TMRCA =15.10Ma (Harvey et al. 2020). Given that bird chromosomes are known to have high synteny and evolutionary stasis between distantly related species (Ellegren 2010), we produced a pseudo-chromosome reference genome for *X. elegans* and *C. ornatus* by ordering and orienting their scaffolds to the chromosomes of the Zebra Finch (*Taeniopygia guttata*; version taeGut3.2.4) with chromosemble in satsuma v3.1.0 (Grabherr et al. 2010). For *R. melanosticta,* we used the chromosome assignment conducted in a previous study (Coelho et al. 2019). To check the completeness of our pseudo-chromosome references, we used Busco v2.0.1 (Waterhouse et al. 2018) to search for a set of single-copy avian ortholog loci. To transfer genome annotations from the scaffold assemblies to the pseudo chromosome reference genomes, we mapped the genomic coordinates of each annotated feature using gmap (Wu and Watanabe 2005). For *R. melanosticta* we used the annotation performed by (Mikkelsen and Weir 2020) and for X*. elegans* and *L. vociferans,* we used the annotations performed by (Feng et al. 2020). A total of 98.90% (15,195), 97.46% (14,834 genes), and 98.92% (15,599 genes) of all annotated genes in *R. melanosticta, X. elegans,* and *C. ornatus* were successfully mapped to the pseudo-chromosome reference, respectively.

We downloaded raw reads from additional closely related species that were used as outgroups in phylogenetic analyses. For *P. nigromaculata,* we included *R. melanosticta, Sakesphorus luctuosus* (GCA_013396695.1 ASM1339669v1; NCBI genome ID: 92896; Feng et al. 2020) and *X. elegans* as outgroups. For *X. spixii,* we included *X. elegans, S. luctuosus, Campylorhamphus procurvoides* (GCA_013396655.1 ASM1339665v1; NCBI genome ID: 92894; Feng et al. 2020), and *Furnarius figulus* (GCA_013397465.1 ASM1339746v1; NCBI genome ID: 92763; Feng et al. 2020). For *L. vociferans,* we included *C. ornatus, Pachyramphus minor* (GCA_013397135.1 ASM1339713v1; NCBI genome ID: 92755; Feng et al. 2020), and *Tyrannus savana* (GCA_013399735.1 ASM1339973v1; NCBI genome ID: 92814; Feng et al. 2020).

### Read alignment, variant calling and filtering

Trimmed and filtered reads were aligned to the references in BWA v0.7.17 (Li and Durbin 2009) using default parameters. We used Picard v.2.0.1 (Broad Institute, Cambridge, MA; http://broadinstitute.github.io/picard/) to 1) sort sam files with SortSam; 2) reassign reads to groups with AddOrReplaceReadGroups; 3) identify duplicated reads with Markduplicates; 4) calculate summary statistics with CollectAlignmentSummaryMetrics, CollectInsertSizeMetrics, and CollectRawWgsMetrics; and 5) create indexes with BuildBamIndex. All Picard functions were run with default parameters. We used the standard GATK v3.8 (McKenna et al. 2010) pipeline to 1) call SNPs and Indels for each individual separately with HaplotypeCaller; 2) perform genotyping with GenotypeGVCFs, assuming a value of 0.05 for the --heterozygosity flag; 3) flag and filter variants with VariantFiltration. Given the lack of a high confidence SNP panel, we implemented hard filtering options recommended by the Broad Institute’s Best Practices (https://gatk.broadinstitute.org/). We filtered SNPs with quality by depth below 2 (QD < 2.0), SNPs where reads containing the alternative allele were considerably shorter than reads with the reference allele (ReadPosRankSum < -8), SNPs with root mean square of the mapping quality lower than 40 (MQ < 40.0), SNPs with evidence of strand bias (FS > 60.0 and SOR > 3.0), and SNPs where the read with the alternative allele had a lower mapping quality than the reference allele (MQRankSumTest < − 12.5). Lastly, we filtered raw VCF files by keeping only bi-allelic sites, with no more than 50% of missing information, with a minimum read depth of 4 and maximum of 30, and read quality score > Q20 using VCFTOOLS v0.1.15 (Danecek et al. 2011). We phased the genotypes in our genomic vcf files using BEAGLE v5.1 (Browning et al.; Browning and Browning 2007) in sliding windows of 10kb and overlap between windows of 1kb.

### Recombination, window-based summary statistics, and genetic structure

To estimate recombination rate (r = recombination rate per base pair per generation) from population level data for each of the species complexes, we used ReLERNN (Adrion et al. 2020). This approach approximates the genomic landscape of recombination by leveraging recurrent neural networks using the raw genotype matrix as a feature vector, avoiding the need to convert the data into summary statistics. ReLERNN calculates r by simulating data matching the θW of the observed DNA sequences.W of the observed DNA sequences. Simulations are then used to train and test a recurrent neural network model designed to predict the per base recombination rate across sliding windows of the genome. Given that genetic structure could potentially influence ReLERNN results (Mezmouk et al. 2011; Mangin et al. 2012), we restricted our analyses to the individuals of the Tapajos interfluve, that was composed exclusively of recent tissue samples, and we did not find any sign of population substructure in the three lineages (see below). Although we did not calculate r for all populations, the landscape of recombination across bird lineages is considered conserved, and variation between recently diverged populations should be minimal (Singhal et al. 2015). To account for the historical demography of the populations, we provided to ReLERNN the output of our SMC++ analyses (see below) with the --demographicHistory option. We considered a mutation rate of 2.42 x 10^-9^ mutations per generation and one year generation time (Jarvis et al. 2014; Zhang et al. 2014).

We calculated population genomics summary statistics for sliding windows using scripts available at https://github.com/simonhmartin/genomics_general. We initially converted vcf files per species into geno format, using parseVCF.py. *Fst*, *Dxy*, and π) across chromosomes for the three studied species. were calculated for the different populations in each of three interfluves using popgenWindows.py. We estimated the D statistics in sliding windows using the ABBABABAwindows.py. We used species tree topology with the highest probability from our species tree analyses (see *Phylogenomic analyses, and topology weighting*) treating the Tapajos, Xingu and Belem populations as the terminals. For all summary statistics, we used phased vcf files, setting the window size to 10kb (-w option) without overlap between windows and the minimum number of sites without missing information per window to 500 (-m option). To obtain GC content proportion across 100kb windows for our reference genomes, we used sequir v4.2 (Charif and Lobry 2007) in R. We fit general linear regressions and Pearson’s correlation index between population genetics summary statistics, phylogenetic weights, and genomic architecture in R. To account for the potential non-linearity of these relationships, we also fit a LOESS model using the R package caret (Kuhn 2008). Models were trained using leave-one-out cross-validation from 80% of the total data.

To explore the genome-wide pattern of genetic structure, we performed Principal Component Analysis (PCA) and individuals relatedness analyses based on identity-by-descent using SNPRelate v1.20.1 (Zheng et al. 2012) in R. In order to minimize the effect of missing genotypes in the PCA, we filtered our vcf files to keep SNPs present in at least 70% of the individuals. We also used SNPRelate to perform an identity-by-state (IBS) analysis among individuals for each species. To avoid the influence of SNP clusters in our PCA and IBS analysis, we pruned SNPs in approximate linkage equilibrium (LD>0.2) with each other.

Specific regions of the genome might be differently affected by selection and gene flow, exhibiting different levels of genetic diversity and differentiation between populations (Langley et al. 2012; Ellegren et al. 2012; Li et al. 2019). To explore the genomic variation in genetic structure, we used lostruct (Li and Ralph 2019). This approach 1) summarizes the relatedness between individuals across genomic windows using PCA, 2) calculates the pairwise dissimilarity in relatedness among window, 3) uses multidimensional scaling (MDS) to produce a visualization of how variable patterns of relatedness are across the genome, and 4) allows the user to combine regions by similarity to inspect contrasting patterns of genetic structure across the genome. We ran lostruct for windows with 1000 SNPs, allowing for 30% of missing genotypes. To visualize the results, we selected the 10% of the windows closer to the three further points on the two first MDS coordinates and performed individual PCA analysis on clustered windows.

### Historical demography, selective sweeps, and linked selection

We modeled population sizes through time using unphased genomes in SMC++ v1.15.3 (Terhorst et al. 2017). Our goal with this approach was to track past fluctuations in *Ne* to be included in ReLERNN (Adrion et al. 2020) and DiploS/HIC (Kern and Schrider 2018) models to account for historical demography. We ran SMC++ exclusively for the Tapajos population of each species assuming a mutation rate of 2.42 x 10^-9^ mutations per generation and one year generation time (Jarvis et al. 2014; Zhang et al. 2014). We explored historical demography of populations within a time window between the present and 300,000ya.

To detect signatures of selection across the genome, we used a Supervised Machine Learning (SML) approach implemented in diploS/HIC (Kern and Schrider 2018). This approach used coalescent simulations of genomic windows to train and test a convolutional neural network (CNN) designed to predict hard and soft selective sweeps and genetic variation linked to selective sweeps across sliding windows of the genome. Genomic windows were simulated using discoal (Kern and Schrider 2016) according to five distinct models: 1) hard selective sweep; 2) soft selective sweep; 3) neutral variation linked to soft selective sweep; 4) neutral variation linked to hard selective sweep; and 5) neutral genetic variation. We performed 5,000 simulations per model using 220kb genomic windows divided into 11 subwindows. To account for the neutral demography of the populations, which is essential to obtain robust model classification between windows (Harris et al. 2018), we added demographic parameters obtained with SMC++ into discoal simulations. To account for uncertainty in simulated parameters, we followed the approach of (Manthey et al. 2021) by allowing current *Ne* to vary between ⅓ to 3x the value obtained with SMC++ within a uniform distribution. Population scaled recombination rate (rho=4Ner; where r is the recombination rate estimated with ReLERNN) priors were set based on the minimum and maximum values obtained across windows with ReLERNN. We set a uniform prior for selection coefficients ranging from 0.00025 to 0.025, and we conditioned sweep completion between the present and 10,000 generations ago. We used a uniform prior between 0.01 and 0.2 for the initial frequency of adaptive variants in soft sweep models. Simulations were converted into feature vectors consisting of population genetics summary statistics, taking into account the observed amount of missing data by using a genomic mask. We calculated the probability of alternative models for observed windows of 20kb. We ran CNNs for 1000 epochs, stopping the run if validation accuracy did not improve for 50 consecutive epochs. We ran five independent runs and predicted observed data with the run that provided the highest accuracy on testing data. To assess the classification power of the CNNs, we inspected the overall accuracy, the false positive rate (FPR), recall (the number of correct positive predictions made out of all positive predictions that could have been made), and area under the curve (AUC). To acknowledge the uncertainty in model selection, we only assigned a model with a probability higher than 0.7 to a genomic window.

### Phylogenomic analyses, and topology weighting

To obtain phylogenetic relationships between individuals, we calculated supermatrix trees concatenating all SNPs using IQTree2 (Minh et al. 2020). We converted vcf files to phylip format using vcf2phylip.py (Ortiz 2019), randomly resolving heterozygous genotypes, and keeping SNPs present in at least 80% of the individuals. In IQTree2 we ran a total of 1000 bootstrap replicates and controlled for ascertainment bias assuming a GTR+ASC substitution model. To obtain phylogenetic trees based on sliding windows of phased vcf files, we used PHYML v3.0 (Guindon et al. 2010) following (Martin and Van Belleghem 2017). We tested windows with different amounts of information content, selecting regions with 50, 100, 500 and 1000 SNPs. We conducted 100 bootstrap replicates per window. To calculate unrooted topology weight for each window across the genome, we used Twisst (Martin and Van Belleghem 2017). This approach allowed us to quantify the relationships among taxa, providing an assessment of the most likely topology for a given genomic region. Given that windows with different information content yielded similar results for the topology weights across the genome, we only present the results for 100 SNPs windows (average window size of 14,503 bp, 15,637 bp, and 5,821 bp for *P. nigromaculata, X. spixii*, and *L. vociferans,* respectively) in subsequent analyses.

To estimate the posterior probability of unrooted species trees, we used Astral-III v5.1.1 (Zhang et al. 2018; Rabiee et al. 2019), using the gene trees produced with phyml as inputs. We used Astral to score unrooted trees (-q option), by calculating their quartet score, branch lengths, and branch support. We set as our main topology (outgroup,Belem(Xingu,Tapajos), and used the -t 2 option to calculate the same metrics for the first alternative and second alternative topologies. Given we only have four terminals per lineage (3 populations + outgroup), three are only three possible unrooted trees. Therefore, this approach allowed us to calculate the posterior probability of all possible topologies. We conducted this approach for the whole set of gene trees and also for subsets of the data, based on specific characteristics of each window. To assess how support for a specific topology varies based on thresholds for specific summary statistics, we selected windows across the genome with the upper and lower 10% tile for recombination rate, Fst, π) across chromosomes for the three studied species., *Dxy* and D statistics.

### Model based approach to account for recombination and selection

In order to explicitly account for gene flow while testing for alternative topologies and estimating demographic parameters of genomic windows, we used a combination of coalescent simulations and supervised machine learning. We simulated data under three alternative topologies, matching the unrooted trees tested in our phylogenetic approach: topology 1) (out,(Belem,(Xingu,Tapajos))); topology 2) (out,(Tapajos,(Xingu,Belem))); and topology 3) (out,(Xingu,(Tapajos,Belem))). We allowed for constant gene flow after the divergence between Xingu and Belem, and Xingu and Tapajos populations. We did not allow gene flow between Belem and Tapajos due to the geographic disjunction between these populations. We simulated 5,000 loci of 10kb, using uniform and wide priors for all parameters (Table S19), and performed one million simulations per model. We assumed a fixed mutation rate of 2.42 x 10^-9^ mutations per generation and a one year generation time (Jarvis et al. 2014; Zhang et al. 2014). Genetic data for each model was simulated in PipeMaster (Gehara et al. 2017), which allows for a user-friendly implementation of msABC (Pavlidis et al. 2010). We summarized genetic variation of observed and simulated data in a feature vector composed of population genetics summary statistics, including mean and variance across loci: number of segregating sites per population and summed across populations; nucleotide diversity per population and for all populations combined; Watterson’s theta (Watterson 1975) per population and for all populations combined; pairwise *Fst* between populations; number of shared alleles between pairs of populations; number of private alleles per population and between pairs of populations; and number of fixed alleles per population and between pairs of populations. To align loci across individuals, phased vcf files per population were split every 10kb windows and converted into a fasta format including monomorphic sites using bcftools (Li 2011). Fasta alignments were converted into feature vectors with PipeMaster which uses PopGenome (Pfeifer et al. 2014) in R. To obtain a genome-wide estimate of demographic parameters, we selected one 10kb genomic window every 100kb to reduce the effect of linkage between windows, and we subsampled 5,000 windows from this data set. We explored how simulated models fitted the observed data PCAs by plotting the first four PCs of simulated statistics vs observed. We also generated goodness-of-fit plots using the gfit function of abc v2.1 (Csilléry et al. 2012) in R.

To classify observed datasets into our three models, we used a neural network (nnet) implemented in Keras v2.3 (https://github.com/rstudio/keras) in R. After an initial exploration for the best architecture for our nnet, we conducted our final analyses using three hidden layers with 32 internal nodes and a “relu” activation function. The output layer was composed of three nodes and a “softmax” activation function. 25% of the simulations were used as testing data. We ran the training step for 1000 epochs using “adam” optimizer and a batch size of 20,000. 5% of the training data set was used for validation, and we used the overall accuray and the sparse_categorical_crossentropy for the loss function to track improvements in model classification. For the most probable model considering genome-wide windows per species, we estimated demographic parameters with a nnet with a similar architecture but designed to predict continuous variables. For this step, we used an output layer with a single node and a “relu” activation. In the training step, we used the mean absolute percentage error (MAE) as an optimizer, training the nnet for 3000 epochs with batch size of 10,000 and a validation split of 0.1. We ran this procedure 10 times for each demographic parameter and summarized the results by calculating the mean across replicates. To additionally assess the accuracy of parameter estimation, we calculated the coefficient of correlation between estimated and true simulated values of the testing data set. To explore how genome-wide parameters differed from regions with distinct signature of selection and under neutrality, we created subsets of 10kb windows that were assigned with high probability (> 0.70) to one of the five distinct models implemented in diploS/HIC. For each species, we estimated parameters based on the best model (topology) considering genome-wide windows. We selected up to 1000 windows for each of the five selection classes and performed the same approach as described above.

To obtain window-based model probability and demographic parameters, we used a similar approach as described above but simulating 100kb window size and using a modified version of PipeMaster (Gehara et al. 2017) that allowed us to simulate intra locus recombination. By selecting a larger window size we increased the information content and resolution of summary statistics of single genomic windows. We performed 100,000 simulations per model, and used the same uniform priors for all parameters as implemented above. For intralocus recombination, we set a uniform prior ranging from 0 to the maximum value obtained with ReLERNN per species (*P. nigromaculata* = 3.021 x 10^-9^; *X. spixii* = 2.475 x 10^-9^; *L. vociferans* = 2.171 x 10^-9^).

Lastly, to explore how recombination rate and gene flow impact topology weight, we performed coalescent simulations based on demographic parameters obtained for *P. nigromaculata,* and calculated topology weights using Twisst (Martin and Van Belleghem 2017). We simulated 1,000 windows of 10kb for four models varying the presence of intra-locus recombination and gene flow between Xingu and Belem, assuming topology 1 (three ingroups plus one outgroup). Simulated parameters are available on Table S20. Simulations were performed with PipeMaster, and we converted the ms output to phylip format with PopGenome. We ran trees for each 10kb window with IQTREE-2 using default parameters and ran Twisst on this estimated set of trees (Figure S15).

## Data access

The raw genetic data underlying this article are available in NCBI short read archive at Bioproject # (accession numbers will be provided upon acceptance). All code and source datasets needed to replicate this study are available at https://doi.org/number will be provided upon acceptance (zenodo):https://github.com/GregoryThom/Genomic-architecture-Amazonian-birds.

## Acknowledgments

We would like to thank K. Provost, J. Merwin, V. Chua, E. Tenorio, J. Cracraft, P. Sweet, T. Trombone, B. Bird, L. Musher. We also thank the Museu Paraense Emílio Goeldi—MPEG for tissue samples. All genetic samples were included on the SisGen platform under the protocols AA7DDBF and AB8BB93. G.T. was funded by the Frank M. Chapman memorial fund of the American Museum of Natural History. Romina Batista received support from Coordenação de Aperfeiçoamento de Pessoal de Nível Superior – Brasil (CAPES-INPA proc. 88887477562/2020-00). B.T.S. was supported by awards from the National Science Foundation US (DEB-1655736; DBI-2029955).

## References

Adrion JR, Galloway JG, Kern AD. 2020. Predicting the Landscape of Recombination Using Deep Learning. Mol Biol Evol 37: 1790–1808.

Albert JS, Craig JM, Tagliacollo VA, Petry P. 2018. Upland and lowland fishes: a test of the river capture hypothesis. Mountains, climate and biodiversity 273–294.

Aleixo A. 2004. Historical diversification of a terra-firme forest bird superspecies: a phylogeographic perspective on the role of different hypotheses of Amazonian diversification. Evolution 58: 1303– 1317.

Aleixo A, Burlamaqui TCT, Schneider MPC. 2009a. Molecular Systematics and Plumage Evolution in the Monotypic Obligate Army-Ant-Following Genus Skutchia (Thamnophilidae). Condor. https://academic.oup.com/condor/article-abstract/111/2/382/5152428.

Aleixo A, Burlamaqui TCT, Schneider MPC, Gonçalves EC. 2009b. Molecular Systematics and Plumage Evolution in the Monotypic Obligate Army-Ant-Following Genus Skutchia (Thamnophilidae)Sistemática Molecular y Evolución del Plumaje en Skutchia, un Género Monotípico que Sigue Ejércitos de Hormigas de Modo Obligatorio (Thamnophilidae). Condor 111: 382–387.

Arbeithuber B, Betancourt AJ, Ebner T, Tiemann-Boege I. 2015. Crossovers are associated with mutation and biased gene conversion at recombination hotspots. Proc Natl Acad Sci U S A 112: 2109–2114.

Barrera-Guzmán AO, Aleixo A, Shawkey MD, Weir JT. 2018. Hybrid speciation leads to novel male secondary sexual ornamentation of an Amazonian bird. Proceedings of the National Academy of Sciences 115: E218–E225.

Barroso GV, Dutheil JY. 2021. Mutation rate variation shapes genome-wide diversity in Drosophila melanogaster. bioRxiv 2021.09.16.460667. https://www.biorxiv.org/content/10.1101/2021.09.16.460667.abstract (Accessed September 23, 2021).

Bates JM, Hackett SJ, Cracraft J. 1998. Area-relationships in the Neotropical lowlands: an hypothesis based on raw distributions of Passerine birds. Journal of Biogeography 25: 783–793. http://dx.doi.org/10.1046/j.1365-2699.1998.2540783.x.

Berv JS, Campagna L, Feo TJ, Castro-Astor I, Ribas CC, Prum RO, Lovette IJ. 2021. Genomic phylogeography of the White-crowned Manakin Pseudopipra pipra (Aves: Pipridae) illuminates a continental-scale radiation out of the Andes. Mol Phylogenet Evol 107205.

Besenbacher S, Hvilsom C, Marques-Bonet T, Mailund T, Schierup MH. 2019. Direct estimation of mutations in great apes reconciles phylogenetic dating. Nat Ecol Evol 3: 286–292.

Bolger AM, Lohse M, Usadel B. 2014. Trimmomatic: a flexible trimmer for Illumina sequence data. Bioinformatics 30: 2114–2120.

Brand CM, White FJ, Ting N, Webster TH. 2021. Soft sweeps predominate recent positive selection in bonobos (Pan paniscus) and chimpanzees (Pan troglodytes). bioRxiv 2020.12.14.422788. https://www.biorxiv.org/content/10.1101/2020.12.14.422788v3.abstract (Accessed June 28, 2021).

Brandvain Y, Kenney AM, Flagel L, Coop G, Sweigart AL. 2014. Speciation and introgression between Mimulus nasutus and Mimulus guttatus. PLoS Genet 10: e1004410.

Bravo GA, Schmitt CJ, Edwards SV. 2021. What Have We Learned from the First 500 Avian Genomes? Annu Rev Ecol Evol Syst 52: 611–639.

Browning BL, Zhou Y, Browning SR. A one penny imputed genome from next generation reference panels. http://dx.doi.org/10.1101/357806.

Browning SR, Browning BL. 2007. Rapid and accurate haplotype phasing and missing-data inference for whole-genome association studies by use of localized haplotype clustering. Am J Hum Genet 81: 1084–1097.

Burri R, Nater A, Kawakami T, Mugal CF, Olason PI, Smeds L, Suh A, Dutoit L, Bureš S, Garamszegi LZ, et al. 2015. Linked selection and recombination rate variation drive the evolution of the genomic landscape of differentiation across the speciation continuum of Ficedula flycatchers. Genome Res 25: 1656–1665.

Byrne H, Lynch Alfaro JW, Sampaio I, Farias I, Schneider H, Hrbek T, Boubli JP. 2018. Titi monkey biogeography: Parallel Pleistocene spread by Plecturocebus and Cheracebus into a post-Pebas Western Amazon. Zool Scr 47: 499–517.

Charif D, Lobry JR. 2007. SeqinR 1.0-2: A Contributed Package to the R Project for Statistical Computing Devoted to Biological Sequences Retrieval and Analysis. In Structural Approaches to Sequence Evolution: Molecules, Networks, Populations (eds. U. Bastolla, M. Porto, H.E. Roman, and M. Vendruscolo), pp. 207–232, Springer Berlin Heidelberg, Berlin, Heidelberg.

Charlesworth B. 1998. Measures of divergence between populations and the effect of forces that reduce variability. Molecular Biology and Evolution 15: 538–543. http://dx.doi.org/10.1093/oxfordjournals.molbev.a025953.

Charlesworth B, Morgan MT, Charlesworth D. 1993. The effect of deleterious mutations on neutral molecular variation. Genetics 134: 1289–1303.

Chase MA, Ellegren H, Mugal CF. 2021. Positive selection plays a major role in shaping signatures of differentiation across the genomic landscape of two independent Ficedula flycatcher species pairs. Evolution 75: 2179–2196.

Coelho LA, Musher LJ, Cracraft J. 2019. A Multireference-Based Whole Genome Assembly for the Obligate Ant-Following Antbird, Rhegmatorhina melanosticta (Thamnophilidae). Diversity 11: 144.

Comeron JM. 2014. Background selection as baseline for nucleotide variation across the Drosophila genome. PLoS Genet 10: e1004434.

Cracraft J. 1985. Historical Biogeography and Patterns of Differentiation within the South American Avifauna: Areas of Endemism. Ornithol Monogr 49–84.

Cruickshank TE, Hahn MW. 2014. Reanalysis suggests that genomic islands of speciation are due to reduced diversity, not reduced gene flow. Mol Ecol 23: 3133–3157.

Csilléry K, François O, Blum MGB. 2012. abc: an R package for approximate Bayesian computation (ABC). Methods Ecol Evol 3: 475–479.

Dagosta FCP, De Pinna M. 2019. The Fishes of the Amazon: Distribution and Biogeographical Patterns, with a Comprehensive List of Species. amnb 2019: 1–163.

Danecek P, Auton A, Abecasis G, Albers CA, Banks E, DePristo MA, Handsaker RE, Lunter G, Marth GT, Sherry ST, et al. 2011. The variant call format and VCFtools. Bioinformatics 27: 2156–2158. http://dx.doi.org/10.1093/bioinformatics/btr330.

Dapper AL, Payseur BA. 2018. Effects of Demographic History on the Detection of Recombination Hotspots from Linkage Disequilibrium. Mol Biol Evol 35: 335–353.

da Silva JMC, Rylands AB, da Fonseca GAB. 2005. The fate of the amazonian areas of endemism. Conserv Biol 19: 689–694.

Delmore KE, Lugo Ramos JS, Van Doren BM, Lundberg M, Bensch S, Irwin DE, Liedvogel M. 2018. Comparative analysis examining patterns of genomic differentiation across multiple episodes of population divergence in birds. Evol Lett 2: 76–87.

Del-Rio G, Rego MA, Whitney BM, Schunck F, Silveira LF, Faircloth BC, Brumfield RT. 2021. Displaced clines in an avian hybrid zone (Thamnophilidae: Rhegmatorhina) within an Amazonian interfluve. Evolution. http://dx.doi.org/10.1111/evo.14377.

de Oliveira DP, de Carvalho VT, Hrbek T. 2016. Cryptic diversity in the lizard genus Plica (Squamata): phylogenetic diversity and Amazonian biogeography. Zool Scr 45: 630–641.

Dutoit L, Burri R, Nater A, Mugal CF, Ellegren H. 2017a. Genomic distribution and estimation of nucleotide diversity in natural populations: perspectives from the collared flycatcher (Ficedula albicollis) genome. Mol Ecol Resour 17: 586–597.

Dutoit L, Vijay N, Mugal CF, Bossu CM, Burri R, Wolf J, Ellegren H. 2017b. Covariation in levels of nucleotide diversity in homologous regions of the avian genome long after completion of lineage sorting. Proceedings of the Royal Society B: Biological Sciences 284. http://dx.doi.org/10.1098/rspb.2016.2756.

Edelman NB, Frandsen PB, Miyagi M, Clavijo B, Davey J, Dikow RB, García-Accinelli G, Van Belleghem SM, Patterson N, Neafsey DE, et al. 2019. Genomic architecture and introgression shape a butterfly radiation. Science 366: 594–599.

Ellegren H. 2010. Evolutionary stasis: the stable chromosomes of birds. Trends Ecol Evol 25: 283–291.

Ellegren H, Smeds L, Burri R, Olason PI, Backström N, Kawakami T, Künstner A, Mäkinen H, Nadachowska-Brzyska K, Qvarnström A, et al. 2012. The genomic landscape of species divergence in Ficedula flycatchers. Nature 491: 756–760.

Elyashiv E, Sattath S, Hu TT, Strutsovsky A, McVicker G, Andolfatto P, Coop G, Sella G. 2016. A Genomic Map of the Effects of Linked Selection in Drosophila. PLoS Genet 12: e1006130.

Ewing GB, Jensen JD. 2016. The consequences of not accounting for background selection in demographic inference. Mol Ecol 25: 135–141.

Farré M, Micheletti D, Ruiz-Herrera A. 2012. Recombination Rates and Genomic Shuffling in Human and Chimpanzee—A New Twist in the Chromosomal Speciation Theory. Mol Biol Evol 30: 853– 864.

Feng S, Stiller J, Deng Y, Armstrong J, Fang Q, Reeve AH, Xie D, Chen G, Guo C, Faircloth BC, et al. 2020. Dense sampling of bird diversity increases power of comparative genomics. Nature 587: 252– 257.

Ferreira M, Fernandes AM, Aleixo A, Antonelli A, Olsson U, Bates JM, Cracraft J, Ribas CC. 2018. Evidence for mtDNA capture in the jacamar Galbula leucogastra/chalcothorax species-complex and insights on the evolution of white-sand ecosystems in the Amazon basin. Mol Phylogenet Evol 129: 149–157.

Fledel-Alon A, Wilson DJ, Broman K, Wen X, Ober C, Coop G, Przeworski M. 2009. Broad-scale recombination patterns underlying proper disjunction in humans. PLoS Genet 5: e1000658.

Fontaine MC, Pease JB, Steele A, Waterhouse RM, Neafsey DE, Sharakhov IV, Jiang X, Hall AB, Catteruccia F, Kakani E, et al. 2015. Mosquito genomics. Extensive introgression in a malaria vector species complex revealed by phylogenomics. Science 347: 1258524.

Garrigan D, Kingan SB, Geneva AJ, Andolfatto P, Clark AG, Thornton KR, Presgraves DC. 2012. Genome sequencing reveals complex speciation in the Drosophila simulans clade. Genome Res 22: 1499–1511.

Gehara M, Garda AA, Werneck FP, Oliveira EF, da Fonseca EM, Camurugi F, Magalhães F de M, Lanna FM, Sites JW Jr, Marques R, et al. 2017. Estimating synchronous demographic changes across populations using hABC and its application for a herpetological community from northeastern Brazil. Mol Ecol 26: 4756–4771.

Gillespie JH. 2000. Genetic drift in an infinite population. The pseudohitchhiking model. Genetics 155: 909–919.

Grabherr MG, Russell P, Meyer M, Mauceli E, Alföldi J, Di Palma F, Lindblad-Toh K. 2010. Genome- wide synteny through highly sensitive sequence alignment: Satsuma. Bioinformatics 26: 1145–1151.

Guindon S, Dufayard J-F, Lefort V, Anisimova M, Hordijk W, Gascuel O. 2010. New algorithms and methods to estimate maximum-likelihood phylogenies: assessing the performance of PhyML 3.0. Syst Biol 59: 307–321.

Haenel Q, Laurentino TG, Roesti M, Berner D. 2018. Meta-analysis of chromosome-scale crossover rate variation in eukaryotes and its significance to evolutionary genomics. Molecular Ecology 27: 2477– 2497. http://dx.doi.org/10.1111/mec.14699.

Haffer J. 2008. Hypotheses to explain the origin of species in Amazonia. Braz J Biol 68: 917–947.

Haffer J. 1969. Speciation in amazonian forest birds. Science 165: 131–137.

Halldorsson BV, Palsson G, Stefansson OA, Jonsson H, Hardarson MT, Eggertsson HP, Gunnarsson B, Oddsson A, Halldorsson GH, Zink F, et al. 2019. Characterizing mutagenic effects of recombination through a sequence-level genetic map. Science 363. http://dx.doi.org/10.1126/science.aau1043.

Harriso RB, Sackman A, Jensen JD. 2018. On the unfounded enthusiasm for soft selective sweeps II: Examining recent evidence from humans, flies, and viruses. PLoS Genet 14: e1007859.

Harvey MG, Bravo GA, Claramunt S, Cuervo AM, Derryberry GE, Battilana J, Seeholzer GF, McKay JS, O’Meara BC, Faircloth BC, et al. 2020. The evolution of a tropical biodiversity hotspot. Science 370: 1343–1348.

Harvey MG, Singhal S, Rabosky DL. 2019. Beyond Reproductive Isolation: Demographic Controls on the Speciation Process. Annu Rev Ecol Evol Syst 50: 75–95.

Hudson RR. 1983. Properties of a neutral allele model with intragenic recombination. Theor Popul Biol 23: 183–201.

Hudson RR, Kaplan NL. 1995. Deleterious background selection with recombination. Genetics 141: 1605–1617. http://dx.doi.org/10.1093/genetics/141.4.1605.

Jarvis ED, Mirarab S, Aberer AJ, Li B, Houde P, Li C, Ho SYW, Faircloth BC, Nabholz B, Howard JT, et al. 2014. Whole-genome analyses resolve early branches in the tree of life of modern birds. Science 346: 1320–1331.

Jensen JD, Payseur BA, Stephan W, Aquadro CF, Lynch M, Charlesworth D, Charlesworth B. 2019. The importance of the Neutral Theory in 1968 and 50 years on: A response to Kern and Hahn 2018. Evolution 73: 111–114.

Johri P, Aquadro CF, Beaumont M, Charlesworth B, Excoffier L, Eyre-Walker A, Keightley PD, Lynch M, McVean G, Payseur BA, et al. Statistical inference in population genomics. http://dx.doi.org/10.1101/2021.10.27.466171.

Johri P, Charlesworth B, Jensen JD. 2020. Toward an Evolutionarily Appropriate Null Model: Jointly Inferring Demography and Purifying Selection. Genetics 215: 173–192.

Johri P, Riall K, Becher H, Excoffier L, Charlesworth B, Jensen JD. 2021. The Impact of Purifying and Background Selection on the Inference of Population History: Problems and Prospects. Mol Biol Evol 38: 2986–3003.

Jónsson H, Sulem P, Arnadottir GA, Pálsson G, Eggertsson HP, Kristmundsdottir S, Zink F, Kehr B, Hjorleifsson KE, Jensson BÖ, et al. 2018. Multiple transmissions of de novo mutations in families. Nat Genet 50: 1674–1680.

Kaback D, Guacci V, Barber D, Mahon J. 1992. Chromosome size-dependent control of meiotic recombination. Science 256: 228–232. http://dx.doi.org/10.1126/science.1566070.

Kartje ME, Jing P, Payseur BA. 2020. Weak Correlation between Nucleotide Variation and Recombination Rate across the House Mouse Genome. Genome Biol Evol 12: 293–299.

Kawakami T, Smeds L, Backström N, Husby A, Qvarnström A, Mugal CF, Olason P, Ellegren H. 2014. A high density linkage map enables a second generation collared flycatcher genome assembly and reveals the patterns of avian recombination rate variation and chromosomal evolution. Molecular Ecology 23: 4035–4058. http://dx.doi.org/10.1111/mec.12810.

Kern AD, Hahn MW. 2018. The Neutral Theory in Light of Natural Selection. Mol Biol Evol 35: 1366– 1371.

Kern AD, Schrider DR. 2018. diploS/HIC: An Updated Approach to Classifying Selective Sweeps. G3 **8**: 1959–1970.

Kern AD, Schrider DR. 2016. Discoal: flexible coalescent simulations with selection. Bioinformatics 32: 3839–3841.

Knowles LL. 2009. Statistical Phylogeography. Annu Rev Ecol Evol Syst 40: 593–612.

Korunes KL, Noor MAF. 2017. Gene conversion and linkage: effects on genome evolution and speciation. Mol Ecol 26: 351–364.

Langley CH, Stevens K, Cardeno C, Lee YCG, Schrider DR, Pool JE, Langley SA, Suarez C, Corbett- Detig RB, Kolaczkowski B, et al. 2012. Genomic variation in natural populations of Drosophila melanogaster. Genetics 192: 533–598.

Li G, Figueiró HV, Eizirik E, Murphy WJ. 2019. Recombination-Aware Phylogenomics Reveals the Structured Genomic Landscape of Hybridizing Cat Species. Mol Biol Evol 36: 2111–2126.

Li H. 2011. A statistical framework for SNP calling, mutation discovery, association mapping and population genetical parameter estimation from sequencing data. Bioinformatics 27: 2987–2993.

Li H, Durbin R. 2009. Fast and accurate short read alignment with Burrows–Wheeler transform. Bioinformatics 25: 1754–1760.

Li H, Ralph P. 2019. Local PCA Shows How the Effect of Population Structure Differs Along the Genome. Genetics 211: 289–304.

Luna LW, Ribas CC, Aleixo A. 2021. Genomic differentiation with gene flow in a widespread Amazonian floodplain specialist bird species. J Biogeogr. https://onlinelibrary.wiley.com/doi/10.1111/jbi.14257.

Lynch Alfaro JW, Boubli JP, Paim FP, Ribas CC, Silva MNF da, Messias MR, Röhe F, Mercês MP, Silva Júnior JS, Silva CR, et al. 2015. Biogeography of squirrel monkeys (genus Saimiri): South-central Amazon origin and rapid pan-Amazonian diversification of a lowland primate. Mol Phylogenet Evol 82 Pt B: 436–454.

Mangin B, Siberchicot A, Nicolas S, Doligez A, This P, Cierco-Ayrolles C. 2012. Novel measures of linkage disequilibrium that correct the bias due to population structure and relatedness. Heredity 108: 285–291.

Manthey JD, Klicka J, Spellman GM. 2021. The genomic signature of allopatric speciation in a songbird is shaped by genome architecture (Aves: Certhia americana). Genome Biol Evol. http://dx.doi.org/10.1093/gbe/evab120.

Martin SH, Davey JW, Salazar C, Jiggins CD. 2019. Recombination rate variation shapes barriers to introgression across butterfly genomes. PLoS Biol 17: e2006288.

Martin SH, Van Belleghem SM. 2017. Exploring Evolutionary Relationships Across the Genome Using Topology Weighting. Genetics 206: 429–438.

McKenna A, Hanna M, Banks E, Sivachenko A, Cibulskis K, Kernytsky A, Garimella K, Altshuler D, Gabriel S, Daly M, et al. 2010. The Genome Analysis Toolkit: a MapReduce framework for analyzing next-generation DNA sequencing data. Genome Res 20: 1297–1303.

McVicker G, Gordon D, Davis C, Green P. 2009. Widespread genomic signatures of natural selection in hominid evolution. PLoS Genet 5: e1000471.

Meunier J, Duret L. 2004. Recombination drives the evolution of GC-content in the human genome. Mol Biol Evol 21: 984–990.

Mezmouk S, Dubreuil P, Bosio M, Décousset L, Charcosset A, Praud S, Mangin B. 2011. Effect of population structure corrections on the results of association mapping tests in complex maize diversity panels. Theor Appl Genet 122: 1149–1160.

Mikkelsen EK, Weir JT. 2020. The genome of the Xingu scale-backed antbird (Willisornis vidua nigrigula) reveals lineage-specific adaptations. Genomics 112: 4552–4560.

Minh BQ, Schmidt HA, Chernomor O, Schrempf D, Woodhams MD, von Haeseler A, Lanfear R. 2020. IQ-TREE 2: New Models and Efficient Methods for Phylogenetic Inference in the Genomic Era. Mol Biol Evol 37: 1530–1534.

Mořkovský L, Janoušek V, Reif J, Rídl J, Pačes J, Choleva L, Janko K, Nachman MW, Reifová R. 2018. Genomic islands of differentiation in two songbird species reveal candidate genes for hybrid female sterility. Mol Ecol 27: 949–958.

Musher LJ, Giakoumis M, Albert J, Del Rio G, Rego M, Thom G, Aleixo A, Ribas CC, Brumfield RT, Smith BT, et al. 2021. River network rearrangements promote speciation in lowland Amazonian birds. bioRxiv 2021.11.15.468717. https://www.biorxiv.org/content/10.1101/2021.11.15.468717v1 (Accessed November 16, 2021).

Nachman MW, Payseur BA. 2012. Recombination rate variation and speciation: theoretical predictions and empirical results from rabbits and mice. Philos Trans R Soc Lond B Biol Sci 367: 409–421.

Nguyen L-T, Schmidt HA, von Haeseler A, Minh BQ. 2015. IQ-TREE: A Fast and Effective Stochastic Algorithm for Estimating Maximum-Likelihood Phylogenies. Molecular Biology and Evolution 32: 268–274. http://dx.doi.org/10.1093/molbev/msu300.

Ortiz EM. 2019. vcf2phylip v2.0: convert a VCF matrix into several matrix formats for phylogenetic analysis. https://zenodo.org/record/2540861.

Pavlidis P, Laurent S, Stephan W. 2010. msABC: a modification of Hudson’s ms to facilitate multi-locus ABC analysis. Mol Ecol Resour 10: 723–727.

Penz C, DeVries P, Tufto J, Lande R. 2015. Butterfly dispersal across Amazonia and its implication for biogeography. Ecography 38: 410–418.

Pessia E, Popa A, Mousset S, Rezvoy C, Duret L, Marais GAB. 2012. Evidence for widespread GC- biased gene conversion in eukaryotes. Genome Biol Evol 4: 675–682.

Pfeifer B, Wittelsbürger U, Ramos-Onsins SE, Lercher MJ. 2014. PopGenome: an efficient Swiss army knife for population genomic analyses in R. Mol Biol Evol 31: 1929–1936.

Pouyet F, Aeschbacher S, Thiéry A, Excoffier L. 2018. Background selection and biased gene conversion affect more than 95% of the human genome and bias demographic inferences. Elife 7. http://dx.doi.org/10.7554/eLife.36317.

Rabiee M, Sayyari E, Mirarab S. 2019. Multi-allele species reconstruction using ASTRAL. Mol Phylogenet Evol 130: 286–296.

Ribas CC, Aleixo A, Nogueira ACR, Miyaki CY, Cracraft J. 2012. A palaeobiogeographic model for biotic diversification within Amazonia over the past three million years. Proc Biol Sci 279: 681–689.

Rousselle M, Mollion M, Nabholz B, Bataillon T, Galtier N. 2018. Overestimation of the adaptive substitution rate in fluctuating populations. Biol Lett 14. http://dx.doi.org/10.1098/rsbl.2018.0055.

Roux C, Fraïsse C, Castric V, Vekemans X, Pogson GH, Bierne N. 2014. Can we continue to neglect genomic variation in introgression rates when inferring the history of speciation? A case study in a Mytilus hybrid zone. J Evol Biol 27: 1662–1675.

Schrider DR, Shanku AG, Kern AD. 2016. Effects of linked selective sweeps on demographic inference and model selection. Genetics 204: 1207–1223.

Schumer M, Xu C, Powell D, Durvasula A, Skov L, Holland C, Sankararaman S, Andolfatto P, Rosenthal G, Przeworski M. 2017. Natural selection interacts with the local recombination rate to shape the evolution of hybrid genomes. bioRxiv: 212407.

Schumer M, Xu C, Powell DL, Durvasula A, Skov L, Holland C, Blazier JC, Sankararaman S, Andolfatto P, Rosenthal GG, et al. 2018. Natural selection interacts with recombination to shape the evolution of hybrid genomes. Science 360: 656–660.

Seehausen O, Butlin RK, Keller I, Wagner CE, Boughman JW, Hohenlohe PA, Peichel CL, Saetre G-P, Bank C, Brännström A, et al. 2014. Genomics and the origin of species. Nat Rev Genet 15: 176–192.

Silva SM, Peterson AT, Carneiro L, Burlamaqui TCT, Ribas CC, Sousa-Neves T, Miranda LS, Fernandes AM, d’Horta FM, Araújo-Silva LE, et al. 2019. A dynamic continental moisture gradient drove Amazonian bird diversification. Sci Adv 5: eaat5752.

Singhal S, Leffler EM, Sannareddy K, Turner I, Venn O, Hooper DM, Strand AI, Li Q, Raney B, Balakrishnan CN, et al. 2015. Stable recombination hotspots in birds. Science 350: 928–932.

Smith BT, McCormack JE, Cuervo AM, Hickerson MJ, Aleixo A, Cadena CD, Pérez-Emán J, Burney CW, Xie X, Harvey MG, et al. 2014. The drivers of tropical speciation. Nature 515: 406–409.

Smith JM, Haigh J. 1974. The hitch-hiking effect of a favourable gene. Genet Res 23: 23–35.

Smith TCA, Arndt PF, Eyre-Walker A. 2018. Large scale variation in the rate of germ-line de novo mutation, base composition, divergence and diversity in humans. PLoS Genet 14: e1007254.

Solís-Lemus C, Ané C. 2016. Inferring Phylogenetic Networks with Maximum Pseudolikelihood under Incomplete Lineage Sorting. PLoS Genet 12: e1005896.

Stankowski S, Chase MA, Fuiten AM, Rodrigues MF, Ralph PL, Streisfeld MA. 2019. Widespread selection and gene flow shape the genomic landscape during a radiation of monkeyflowers. http://dx.doi.org/10.1101/342352.

Terhorst J, Kamm JA, Song YS. 2017. Robust and scalable inference of population history from hundreds of unphased whole genomes. Nat Genet 49: 303–309.

Tigano A, Khan R, Omer AD, Weisz D, Dudchenko O, Multani AS, Pathak S, Behringer RR, Aiden EL, Fisher H, et al. 2021. Chromosome size affects sequence divergence between species through the interplay of recombination and selection. bioRxiv 2021.01.15.426870. https://www.biorxiv.org/content/10.1101/2021.01.15.426870v1 (Accessed June 7, 2021).

Van Belleghem SM, Baquero M, Papa R, Salazar C, McMillan WO, Counterman BA, Jiggins CD, Martin SH. 2018. Patterns of Z chromosome divergence among Heliconius species highlight the importance of historical demography. Mol Ecol 27: 3852–3872.

Van Doren BM, Campagna L, Helm B, Illera JC, Lovette IJ, Liedvogel M. 2017. Correlated patterns of genetic diversity and differentiation across an avian family. Mol Ecol 26: 3982–3997.

Vijay N, Weissensteiner M, Burri R, Kawakami T, Ellegren H, Wolf JBW. 2017. Genomewide patterns of variation in genetic diversity are shared among populations, species and higher-order taxa. Mol Ecol 26: 4284–4295.

Waterhouse RM, Seppey M, Simão FA, Manni M, Ioannidis P, Klioutchnikov G, Kriventseva EV, Zdobnov EM. 2018. BUSCO Applications from Quality Assessments to Gene Prediction and Phylogenomics. Mol Biol Evol 35: 543–548.

Watterson GA. 1975. On the number of segregating sites in genetical models without recombination. Theor Popul Biol 7: 256–276.

Wayne ML, Simonsen KL. 1998. Statistical tests of neutrality in the age of weak selection. Trends Ecol Evol 13: 236–240.

Weir JT, Faccio MS, Pulido-Santacruz P, Barrera-Guzmán AO, Aleixo A. 2015. Hybridization in headwater regions, and the role of rivers as drivers of speciation in Amazonian birds. Evolution 69: 1823–1834.

Wen D, Yu Y, Zhu J, Nakhleh L. 2018. Inferring Phylogenetic Networks Using PhyloNet. Syst Biol 67: 735–740.

Wolf JBW, Ellegren H. 2017. Making sense of genomic islands of differentiation in light of speciation. Nat Rev Genet 18: 87–100.

Wu TD, Watanabe CK. 2005. GMAP: a genomic mapping and alignment program for mRNA and EST sequences. Bioinformatics 21: 1859–1875.

Zeng K. 2013. A coalescent model of background selection with recombination, demography and variation in selection coefficients. Heredity 110: 363–371. http://dx.doi.org/10.1038/hdy.2012.102.

Zhang C, Rabiee M, Sayyari E, Mirarab S. 2018. ASTRAL-III: polynomial time species tree reconstruction from partially resolved gene trees. BMC Bioinformatics 19: 153.

Zhang G, Li C, Li Q, Li B, Larkin DM, Lee C, Storz JF, Antunes A, Greenwold MJ, Meredith RW, et al. 2014. Comparative genomics reveals insights into avian genome evolution and adaptation. Science 346: 1311–1320.

Zheng X, Levine D, Shen J, Gogarten SM, Laurie C, Weir BS. 2012. A high-performance computing toolset for relatedness and principal component analysis of SNP data. Bioinformatics 28: 3326–3328.

Edwards SV, Robin V, Ferrand N, Moritz C. 2021. The evolution of comparative phylogeography: putting the geography (and more) into comparative population genomics. Genome Biol. Evol.: http://dx.doi.org/10.1093/gbe/evab176

Kuhn M. 2008. Building predictive models in R using the caret package. J. Stat. Softw. 28, 1–26.

Thom G, Aleixo A. 2015. Cryptic speciation in the white-shouldered antshrike (Thamnophilus aethiops, Aves – Thamnophilidae): The tale of a transcontinental radiation across rivers in lowland Amazonia and the northeastern Atlantic Forest. Molecular Phylogenetics and Evolution 82:95–110.

Thom G, Xue AT, Sawakuchi AO, Ribas CC, Hickerson MJ, Aleixo A, Miyaki C. 2020. Quaternary climate changes as speciation drivers in the Amazon floodplains. Sci Adv 6:eaax4718.

