## Supplementary Material for "Genomic architecture drives population structuring in Amazonian birds" (from another version of this preprint: version 1, 2021-12-02)

**Supplementary information**

**Suplementary Tables**

Table S1:Specimens sequenced in this study for three species of Amazonian Birds. Available at https://github.com/GregoryThom/Genomic-architecture-Amazonian-birds (DOI will be provided upon acceptance)

Tables S2: Benchmarking Universal Single-Copy Orthologs analyses (BUSCO) output.

| | R. melanosticta | X. elegans | C. ornatus |
|---|---|---|---|
| Complete BUSCOs (C) | 4389 | 4378 | 4588 |
| Complete and single-copy BUSCOs (S) | 4341 | 4320 | 4536 |
| Complete and duplicated BUSCOs (D) | 48 | 58 | 52 |
| Fragmented BUSCOs (F) | 163 | 272 | 209 |
| Missing BUSCOs (M) | 363 | 265 | 118 |
| Total BUSCO groups searched | 4915 | 4915 | 4915 |
| Total % | 89.3 | 89.1 | 93.4 |

Table S3: Summary statistics calculated for 100kb non overlapping widows for *Phlegopsis nigromaculata*. Available at https://github.com/GregoryThom/Genomic-architecture-Amazonian-birds (DOI will be provided upon acceptance)

Table S4: Summary statistics calculated for 100kb non overlapping widows for *Xiphorhynchus spixii*. Available at https://github.com/GregoryThom/Genomic-architecture-Amazonian-birds (DOI will be provided upon acceptance)

Table S5: Summary statistics calculated for 100kb non overlapping widows for *Lipaugus vociferans*. Available at https://github.com/GregoryThom/Genomic-architecture-Amazonian-birds (DOI will be provided upon acceptance)

Table S6: P-values for the correlogram (Pearson's correlation index) between population genetics summary statistics, phylogenetic weights and genomic architecture estimates for *Phlegopsis nigromaculata*. Correlogram is available on Figure S1. Available at https://github.com/GregoryThom/Genomic-architecture-Amazonian-birds (DOI will be provided upon acceptance)

Table S7: P-values for the correlogram (Pearson's correlation index) between population genetics summary statistics, phylogenetic weights and genomic architecture estimates for *Xiphorhynchus spixii*. Correlogram is available on Figure S2. Available at https://github.com/GregoryThom/Genomic-architecture-Amazonian-birds (DOI will be provided upon acceptance)

Table S8: P-values for the correlogram (Pearson's correlation index) between population genetics summary statistics, phylogenetic weights and genomic architecture estimates for *Lipaugus vociferans* Correlogram is available on Figure S3. Available at https://github.com/GregoryThom/Genomic-architecture-Amazonian-birds (DOI will be provided upon acceptance)

Table S9: P-values for the correlogram (Pearson's correlation index) between population genetics summary statistics, phylogenetic weights and genomic architecture estimates for *Phlegopsis nigromaculata*, based on the average values per chromosome. Correlogram is available on Figure S4. Available at https://github.com/GregoryThom/Genomic-architecture-Amazonian-birds (DOI will be provided upon acceptance)

Table S10: P-values for the correlogram (Pearson's correlation index) between population genetics summary statistics, phylogenetic weights and genomic architecture estimates for *Xiphorhynchus spixii*, based on the average values per chromosome. Correlogram is available on Figure S5. Available at https://github.com/GregoryThom/Genomic-architecture-Amazonian-birds (DOI will be provided upon acceptance)

Table S11: P-values for the correlogram (Pearson's correlation index) between population genetics summary statistics, phylogenetic weights and genomic architecture estimates for *Lipaugus vociferans*, based on the average values per chromosome. Correlogram is available on Figure S6. Available at https://github.com/GregoryThom/Genomic-architecture-Amazonian-birds (DOI will be provided upon acceptance)

Table S12: LOESS models results between population genetics summary statistics and genomic architecture variables.

| Species | Model | span | RMSE | $R^2$ | MAE |
|---|---|---|---|---|---|
| *P. nigromaculata* | Topology 1 weight ~ Recombination | 0.5 | 986.341 | 0.040 | 741.486 |

| | | | | | |
|---|---|---|---|---|---|
| *P. nigromaculata* | Topology 2 weight ~ Recombination | 0.5 | 1019.543 | 0.064 | 770.654 |
| *P. nigromaculata* | Topology 3 weight ~ Recombination | 0.5 | 895.657 | 0.010 | 671.060 |
| *P. nigromaculata* | D statistics ~ Recombination | 0.5 | 0.105 | 0.030 | 0.079 |
| *P. nigromaculata* | Theta Pi Tapajos ~ Recombination | 0.5 | 0.001 | 0.325 | 0.001 |
| *P. nigromaculata* | Theta Pi Tapajos ~ Recombination + Gene density | 0.5 | 0.001 | 0.329 | 0.001 |
| *P. nigromaculata* | Gene density ~ Recombination | 0.5 | 0.030 | 0.023 | 0.022 |
| *P. nigromaculata* | Theta Pi Tapajos ~ GC content | 0.5 | 0.001 | 0.110 | 0.001 |
| *X. spixii* | Topology 1 weight ~ Recombination | 0.5 | 641.862 | 0.113 | 468.449 |
| *X. spixii* | Topology 2 weight ~ Recombination | 0.5 | 1145.018 | 0.147 | 823.209 |
| *X. spixii* | Topology 3 weight ~ Recombination | 0.5 | 630.543 | 0.134 | 455.687 |
| *X. spixii* | D statistics ~ Recombination | 0.5 | 0.059 | 0.004 | 0.044 |
| *X. spixii* | Theta Pi Tapajos ~ Recombination | 0.5 | 0.002 | 0.431 | 0.001 |
| *X. spixii* | Theta Pi Tapajos ~ Recombination + Gene density | 0.5 | 0.001 | 0.650 | 0.001 |
| *X. spixii* | Gene density ~ Recombination | 0.5 | 0.023 | 0.030 | 0.017 |

| | | | | | |
|---|---|---|---|---|---|
| *X. spixii* | Theta Pi Tapajos ~ GC content | 0.5 | 0.002 | 0.260 | 0.001 |
| *L. vociferans* | Topology 1 weight ~ Recombination | 0.5 | 113.203 | 0.004 | 87.709 |
| *L. vociferans* | Topology 2 weight ~ Recombination | 0.5 | 134.331 | 0.027 | 105.043 |
| *L. vociferans* | Topology 3 weight ~ Recombination | 0.5 | 136.926 | 0.029 | 105.716 |
| *L. vociferans* | D statistics ~ Recombination | 0.5 | 0.089 | 0.024 | 0.066 |
| *L. vociferans* | Theta Pi Tapajos ~ Recombination | 0.5 | 0.001 | 0.353 | 0.001 |
| *L. vociferans* | Theta Pi Tapajos ~ Recombination + Gene density | 0.5 | 0.001 | 0.406 | 0.001 |
| *L. vociferans* | Gene density ~ Recombination | 0.5 | 0.022 | 0.027 | 0.013 |
| *L. vociferans* | Theta Pi Tapajos ~ GC content | 0.5 | 0.001 | 0.031 | 0.001 |

Table S13: Neural Network testing step performance for the multi-class model classification implemented in DiploS/HIC.

| Parameter | *P. nigromaculata* | *L. vociferans* | *X. spixii* |
|---|---|---|---|
| Loss | 0.924 | 0.856 | 0.781 |
| Accuracy | 0.722 | 0.686 | 0.688 |
| FPR | 0.258 | 0.276 | 0.278 |
| Recall | 0.698 | 0.662 | 0.666 |
| AUC | 0.939 | 0.92 | 0.927 |

FPR - False Positive Rate; AUC - Area Under the Curve

Table S14: Demographic parameter estimation for *P. nigromaculata* obtained with a Neural Network regression model based on 5,000 genomic windows of 10kb.

| Parameter | Average across 10 replicates | SD | $R^2$ | MAE |
|---|---|---|---|---|
| Ne Tapajos | 259924.184 | 12977.39 | 0.993 | 12801.096 |
| Ne Xingu | 169100.342 | 6053.634 | 0.999 | 6531.002 |
| Ne Belem | 332015.849 | 21924.035 | 0.994 | 12469.778 |
| Divergence time Tapajos/Xingu | 77866.978 | 17849.052 | 0.969 | 37810.559 |
| Divergence time Belem(Tapajos/Xingu) | 149836.929 | 15272.708 | 0.974 | 36576.586 |
| Migration Tapajos -> Xingu | 0.002 | 0.005 | 0.917 | 0.161 |
| Migration Tapajos <- Xingu | 0.881 | 0.659 | 0.776 | 0.301 |
| Migration Xingu -> Belem | 0.484 | 0.486 | 0.962 | 0.138 |
| Migration Xingu -> Belem | 0.035 | 0.072 | 0.861 | 0.198 |

Ne - Effective population size

Table S15: Demographic parameter estimation for *X. spixii* obtained with a Neural Network regression model based on 5,000 genomic windows of 10kb.

| Parameters | Average across 10 replicates | SD | $R^2$ | MAE |
|---|---|---|---|---|
| Ne Tapajos | 517153.561 | 34287.162 | 0.995 | 11445.349 |
| Ne Xingu | 488411.928 | 64266.264 | 0.995 | 11262.503 |
| Ne Belem | 168861.451 | 28343.54 | 0.998 | 6569.784 |

| | | | | |
|---|---|---|---|---|
| | | 1 | | |
| Divergence time Tapajos/Xingu | 40303.463 | 16236.143 | 0.973 | 32798.519 |
| Divergence time Belem(Tapajos/Xingu) | 218858.047 | 12095.599 | 0.986 | 30668.129 |
| Migration Tapajos -> Xingu | 0.965 | 0.662 | 0.879 | 0.182 |
| Migration Tapajos <- Xingu | 0.745 | 0.145 | 0.972 | 0.106 |
| Migration Xingu -> Belem | 1.25 | 0.383 | 0.895 | 0.186 |
| Migration Xingu -> Belem | 2.075 | 0.144 | 0.939 | 0.139 |

Ne - Effective population size

Table S16: Demographic parameter estimation for *L. vociferans* obtained with a Neural Network regression model based on 5,000 genomic windows of 10kb.

| Parameters | Average across 10 replicates | $R^2$ | MAE |
|---|---|---|---|
| Ne Tapajos | 395748.563 | 0.983 | 13956.26 |
| Ne Xingu | 409559.188 | 0.98 | 7546.477 |
| Ne Belem | 294531.406 | 0.97 | 12449.989 |
| Divergence time Tapajos/Xingu | 26090.762 | 0.87 | 3382.221 |
| Divergence time Belem(Tapajos/Xingu) | 36558.706 | 0.874 | 2728.545 |
| Migration Tapajos -> Xingu | 2.347 | 0.852 | 0.175 |
| Migration Tapajos <- Xingu | 1.956 | 0.792 | 0.225 |
| Migration Xingu -> Belem | 1.827 | 0.863 | 0.205 |
| Migration Xingu -> Belem | 2.125 | 0.923 | 0.179 |

Ne - Effective population size

Table S17: Model probability for the three alternative unrooted topologies between Tapajos, Xingu, and Belem areas of endemism (plus outgroup), obtained with a Neural Network multi-class model classification approach based on 5,000 genomic windows of 10kb. ______

| Species | Topology 1 | Topology 2 | Topology 3 |
|---|---|---|---|
| *Phlegopsis nigromaculata* | 0.865 | 0.124 | 0.011 |
| *Xiphorhynchus spixii* | 0.000 | 1.000 | 0.000 |
| *Lipaugus vociferans* | 0.542 | 0.285 | 0.172 |

Table S18: LOESS models for demographic parameters estimated with a Neural Network regression model based on genome-wide windows of 100kb as response variables, and recombination rate as predictor variable for the three studied species.

| Species | span | RMSE | $R^2$ | MAE | Response variable |
|---|---|---|---|---|---|
| *P. nigromaculata* | 0.5 | 0.4527 | 0.0055 | 0.4336 | Probability for Topology 1 |
| *P. nigromaculata* | 0.5 | 0.4527 | 0.0049 | 0.4335 | Probability for Topology 2 |
| *P. nigromaculata* | 0.5 | 88313.7500 | 0.2725 | 69768.1481 | Ne Tapajos |
| *P. nigromaculata* | 0.5 | 0.1086 | 0.1405 | 0.0873 | Gene flow |
| *P. nigromaculata* | 0.5 | 28463.0250 | 0.0172 | 21342.1566 | TMRCA |
| *X. spixii* | 0.5 | 0.0537 | 0.0028 | 0.0087 | Probability for Topology 1 |
| *X. spixii* | 0.5 | 0.0526 | 0.0037 | 0.0087 | Probability for Topology 2 |

| | | | | | |
|---|---|---|---|---|---|
| *X. spixii* | 0.5 | 93266.0627 | 0.5648 | 72882.1395 | Ne Tapajos |
| *X. spixii* | 0.5 | 0.0823 | 0.2621 | 0.0661 | Gene flow |
| *X. spixii* | 0.5 | 46318.1524 | 0.0086 | 34985.3272 | TMRCA |
| *L. vociferans* | 0.5 | 0.1109 | 0.1520 | 0.0390 | Probability for Topology 1 |
| *L. vociferans* | 0.5 | 0.1105 | 0.1502 | 0.0390 | Probability for Topology 2 |
| *L. vociferans* | 0.5 | 73405.4776 | 0.3645 | 57788.2004 | Ne Tapajos |
| *L. vociferans* | 0.5 | 0.0246 | 0.0242 | 0.0186 | Gene flow |
| *L. vociferans* | 0.5 | 8567.5522 | 0.1832 | 6628.1080 | TMRCA |

Ne - Effective population size; TMRCA - Time to the most recent common ancestor of the most recent divergence event; Gene flow - Gene flow rate between Tapajos and Xingu populations.

Table S19: Set of uniform priors for demographic parameters implemented in our Neural network approaches.

| Parameter | Lower bound | Upper bound |
|---|---|---|
| Ne Tapajos | 50,000 | 700,000 |
| Ne Xingu | 50,000 | 700,000 |
| Ne Belem | 50,000 | 700,000 |
| Migration Tapajos -> Xingu | 0.01 | 2.00 |
| Migration Tapajos -> Belem | 0.00 | 0.00 |
| Migration Xingu -> Tapajos | 0.01 | 2.00 |
| Migration Xingu -> Belem | 0.01 | 2.00 |
| Migration Belem -> Tapajos | 0.00 | 0.00 |
| Migration Belem -> Xingu | 0.01 | 2.00 |

| | | |
|---|---|---|
| Divergence Time (generations) Tapajos/Xingu | 10,000 | 1,000,000 |
| Divergence Time (generations) Belem(Tapajos/Xingu) | 10,000 | 1,000,000 |
| Mutation rate (u) | 2.42E-09 | 2.42E-09 |

Ne - Effective population size.

Table S20: Demographic parameters used in our simulation approach to explore the effects of recombination and gene flow on topology weights. We tested 4 alternative models varying the presence of intra-locus recombination and gene flow among non sister taxa.

| Parameters | No recombination/No gene flow | Recombination/ No gene flow | No recombination/Gene flow | Recombination/ Gene flow |
|---|---|---|---|---|
| Ne pop1 | 20000 | 20000 | 20000 | 20000 |
| Ne pop2 | 20000 | 20000 | 20000 | 20000 |
| Ne pop3 | 20000 | 20000 | 20000 | 20000 |
| Ne outgroup | 20000 | 20000 | 20000 | 20000 |
| Tdiv 1-2 | 100000 | 100000 | 100000 | 100000 |
| Tdiv 3(1-2) | 300000 | 300000 | 300000 | 300000 |
| Tdiv 4(3(1-2)) | 700000 | 700000 | 700000 | 700000 |
| mig 2->3 | 0 | 0 | 1-10 | 1-10 |
| mig 3->2 | 0 | 0 | 1-10 | 1-10 |
| Recombination | 0 | 1E-08 | 0 | 1E-08 |
| Mutation Rate | 2.42E-09 | 2.42E-09 | 2.42E-09 | 2.42E-09 |

Ne - Effective population size; Tdiv - Divergence time; mig - gene flow.

### **Supplementary Figures**

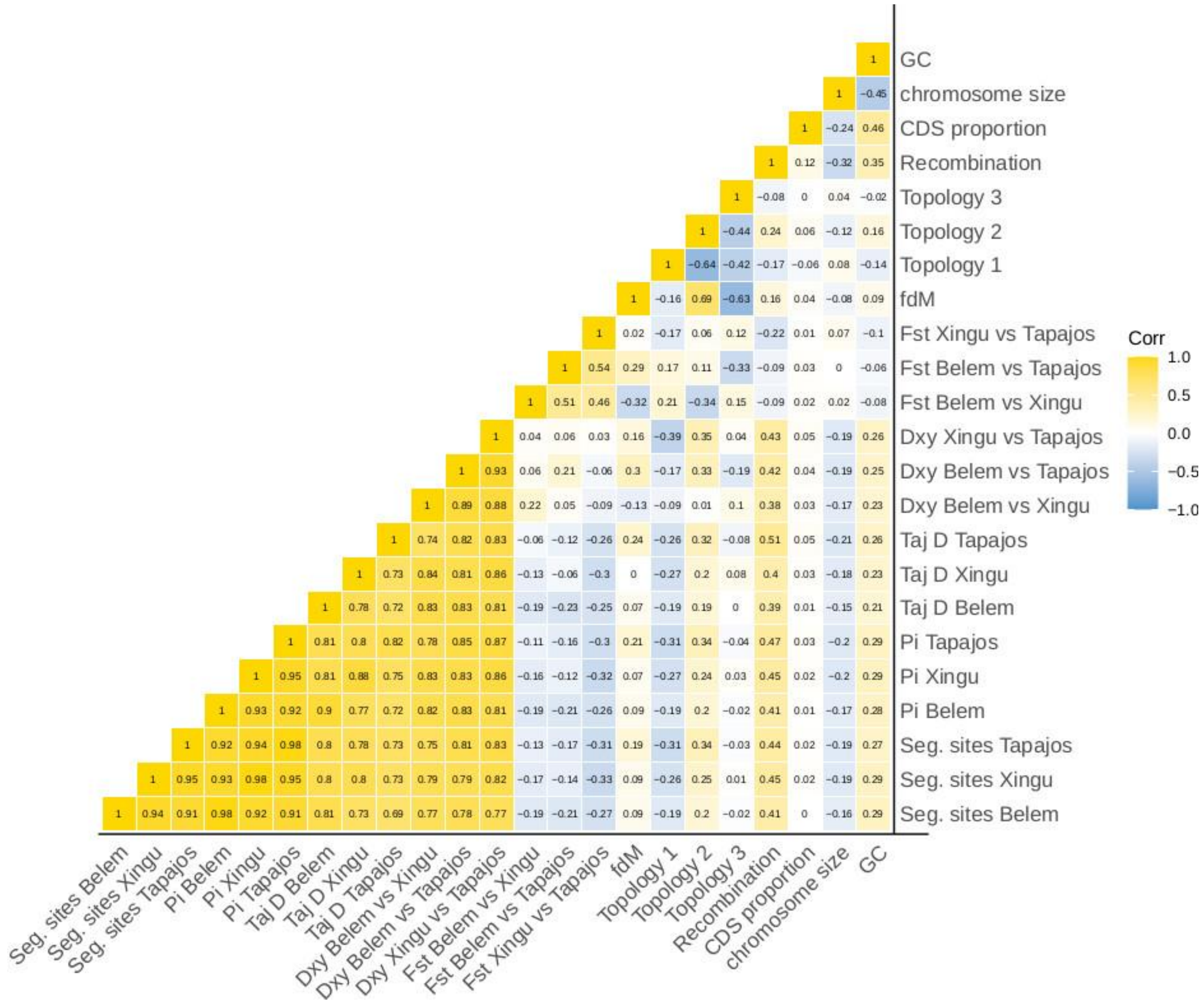


Figure S1: Correlogram (Pearson's correlation index) between population genetics summary statistics, phylogenetic weights and genomic architecture estimates for *Phlegopsis nigromaculata*. P-values are available on supplementary table 6.

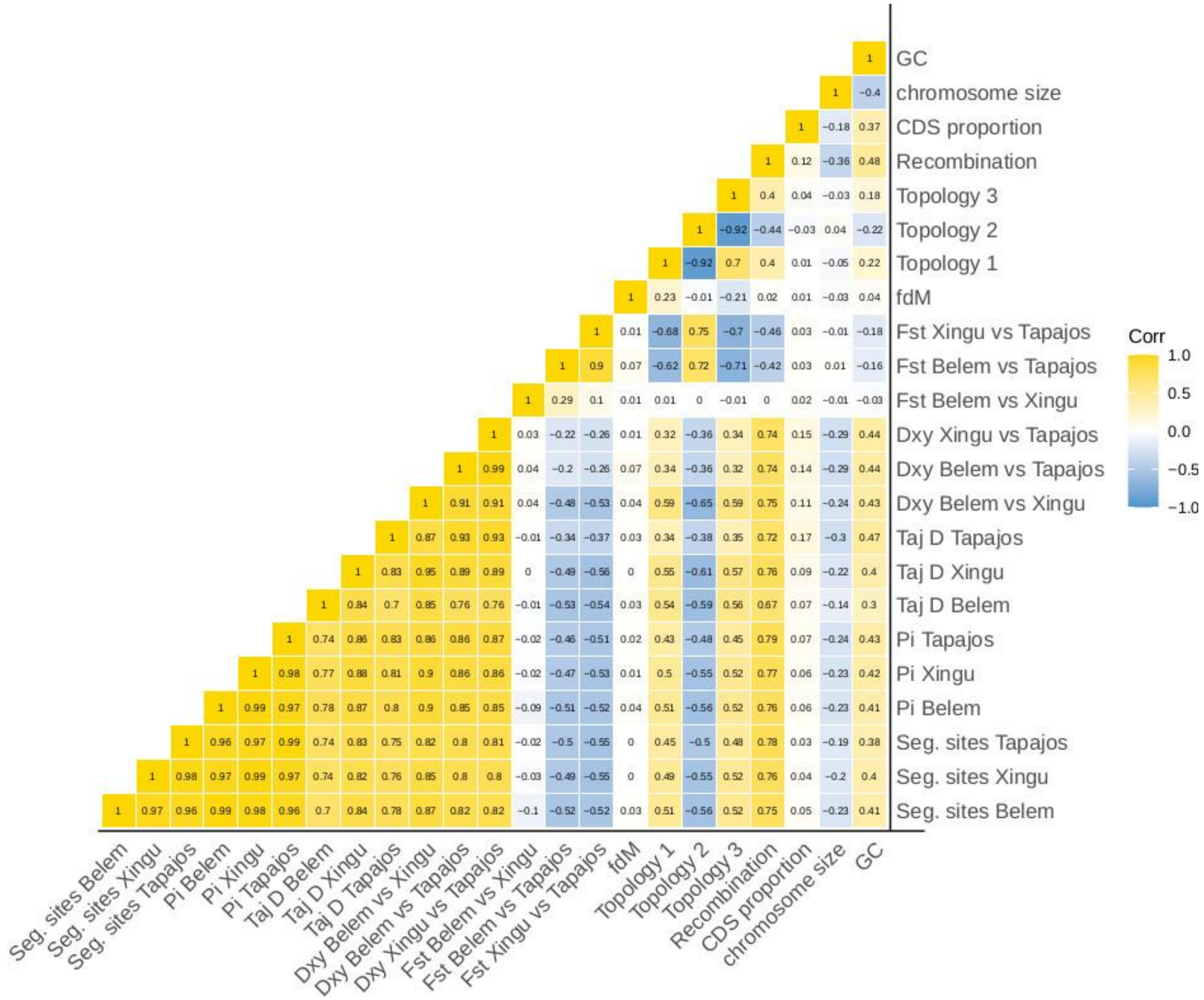


Figure S2: Correlogram (Pearson's correlation index) between population genetics summary statistics, phylogenetic weights and genomic architecture estimates for *Xiphorhynchus spixii*. P-values are available on supplementary table 7.

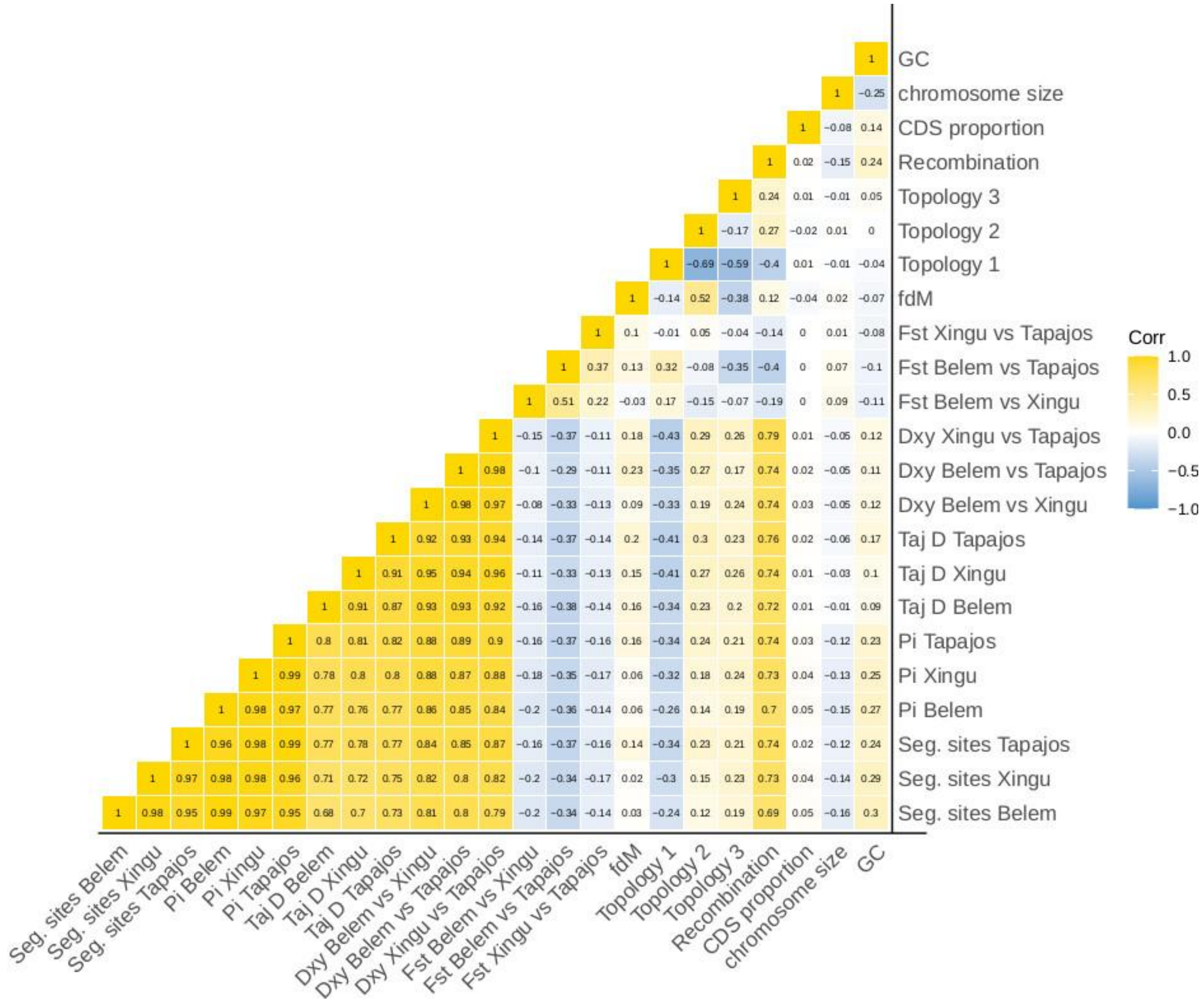


Figure S3: Correlogram (Pearson's correlation index) between population genetics summary statistics, phylogenetic weights and genomic architecture estimates for *Lipaugus vociferans*. P-values are available on supplementary table 8.

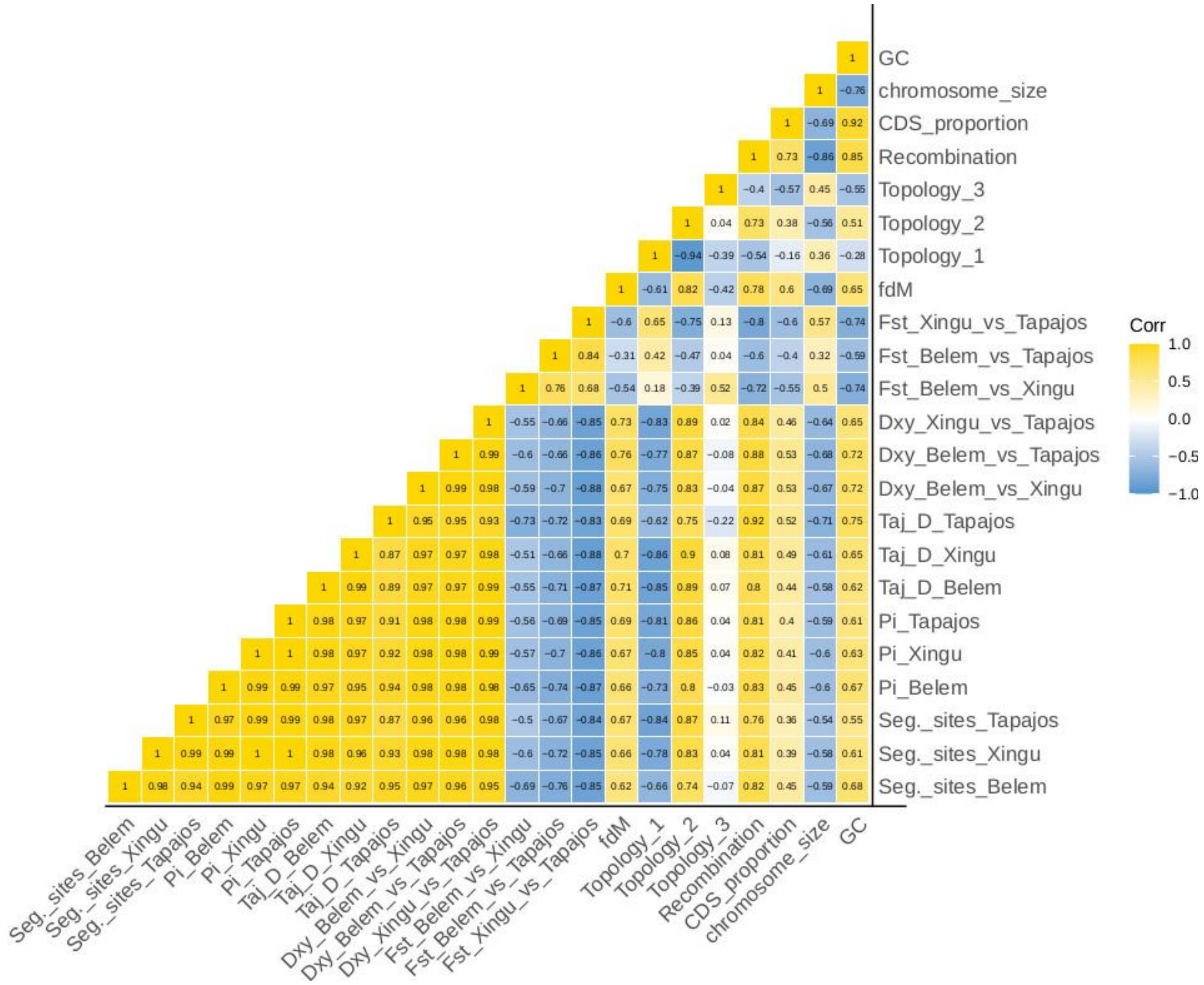


Figure S4: Correlogram (Pearson's correlation index) between population genetics summary statistics, phylogenetic weights and genomic architecture estimates for *P. nigromaculata*, based on the average values per chromosome. P-values are available on supplementary table 9.

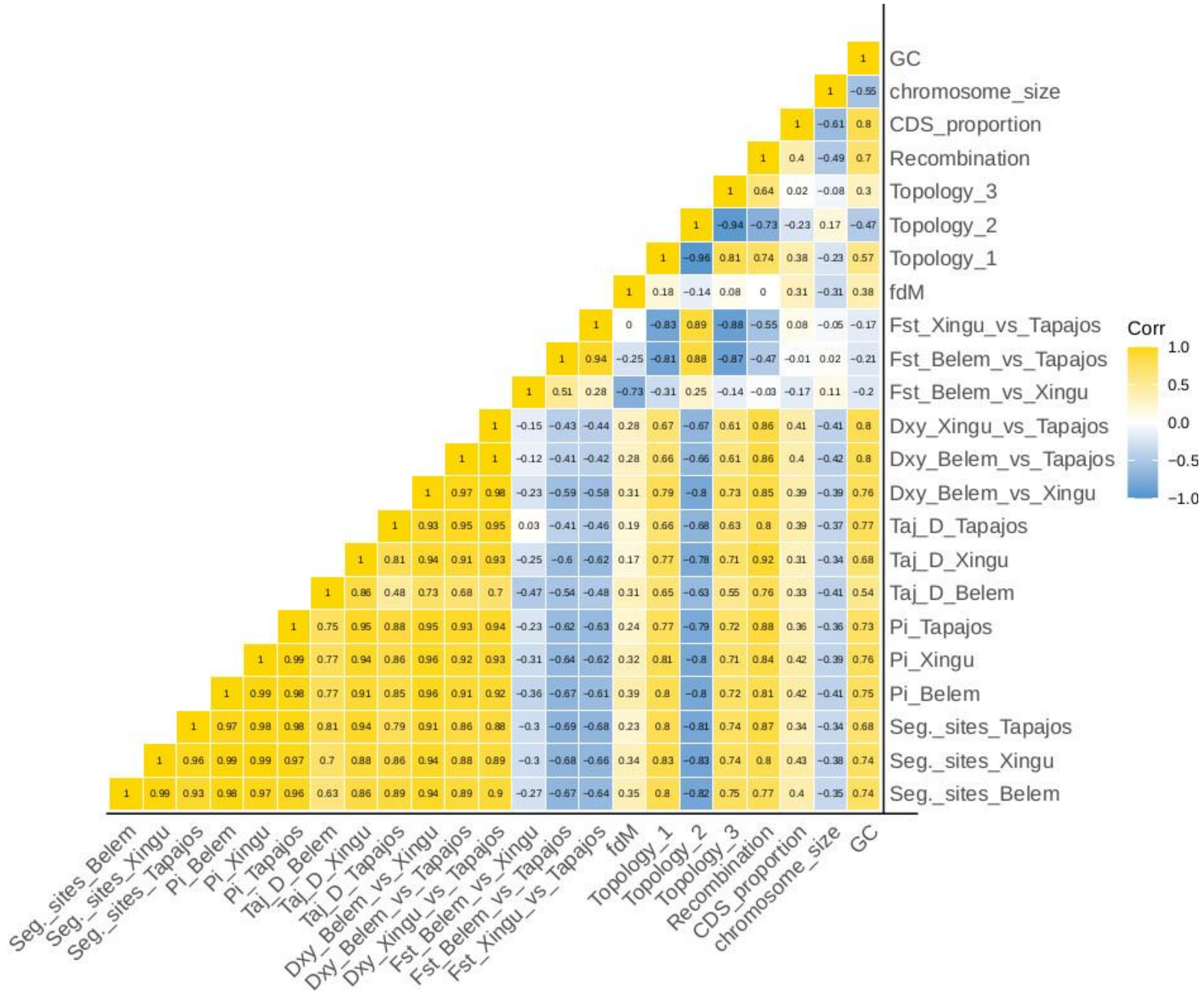


Figure S5: Correlogram (Pearson's correlation index) between population genetics summary statistics, phylogenetic weights and genomic architecture estimates for *Xiphorhynchus spixii*, based on the average values per chromosome. P-values are available on supplementary table 10.

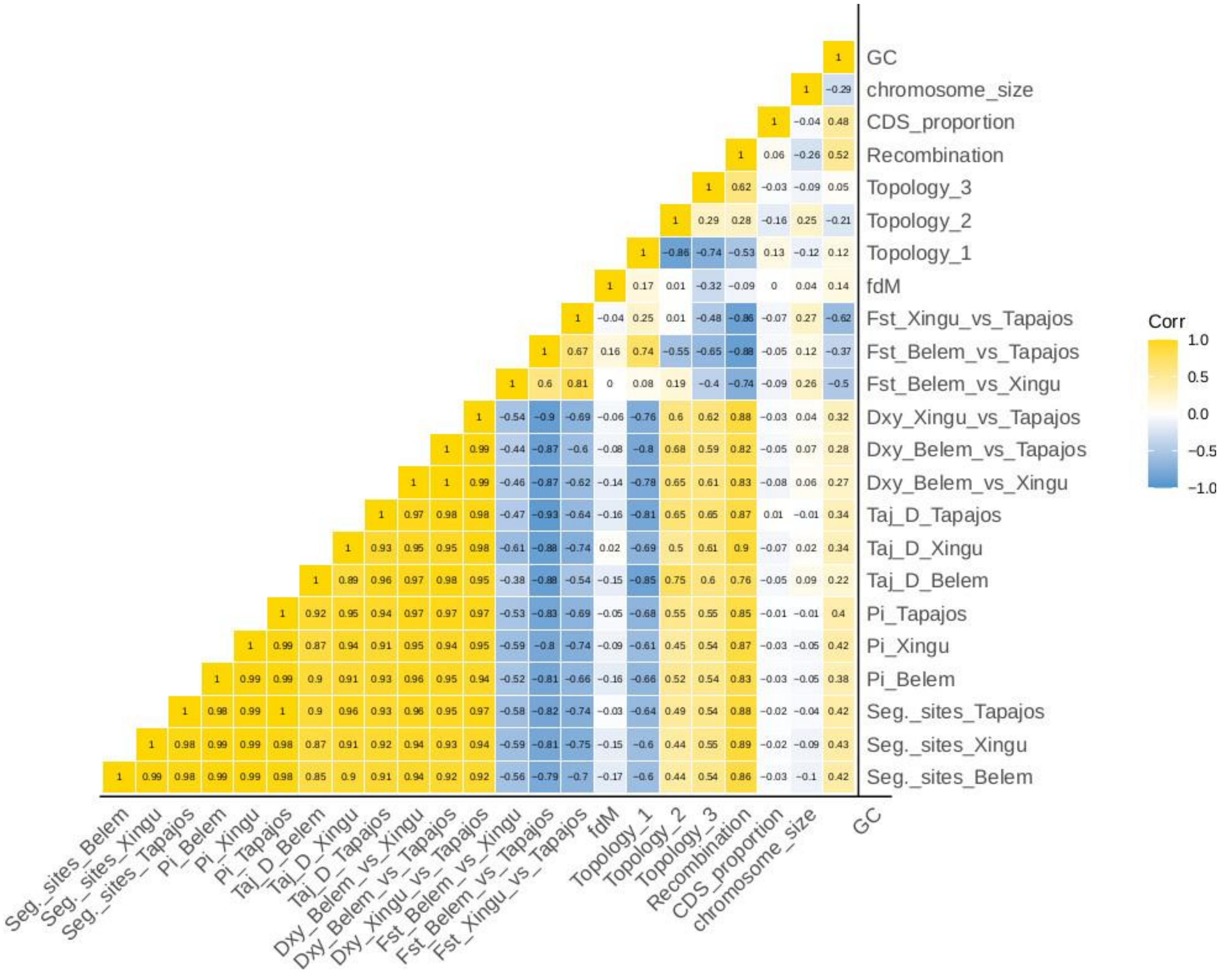


Figure S6: Correlogram (Pearson's correlation index) between population genetics summary statistics, phylogenetic weights and genomic architecture estimates for *Lipaugus vociferans*, based on the average values per chromosome. P-values are available on supplementary table 11.

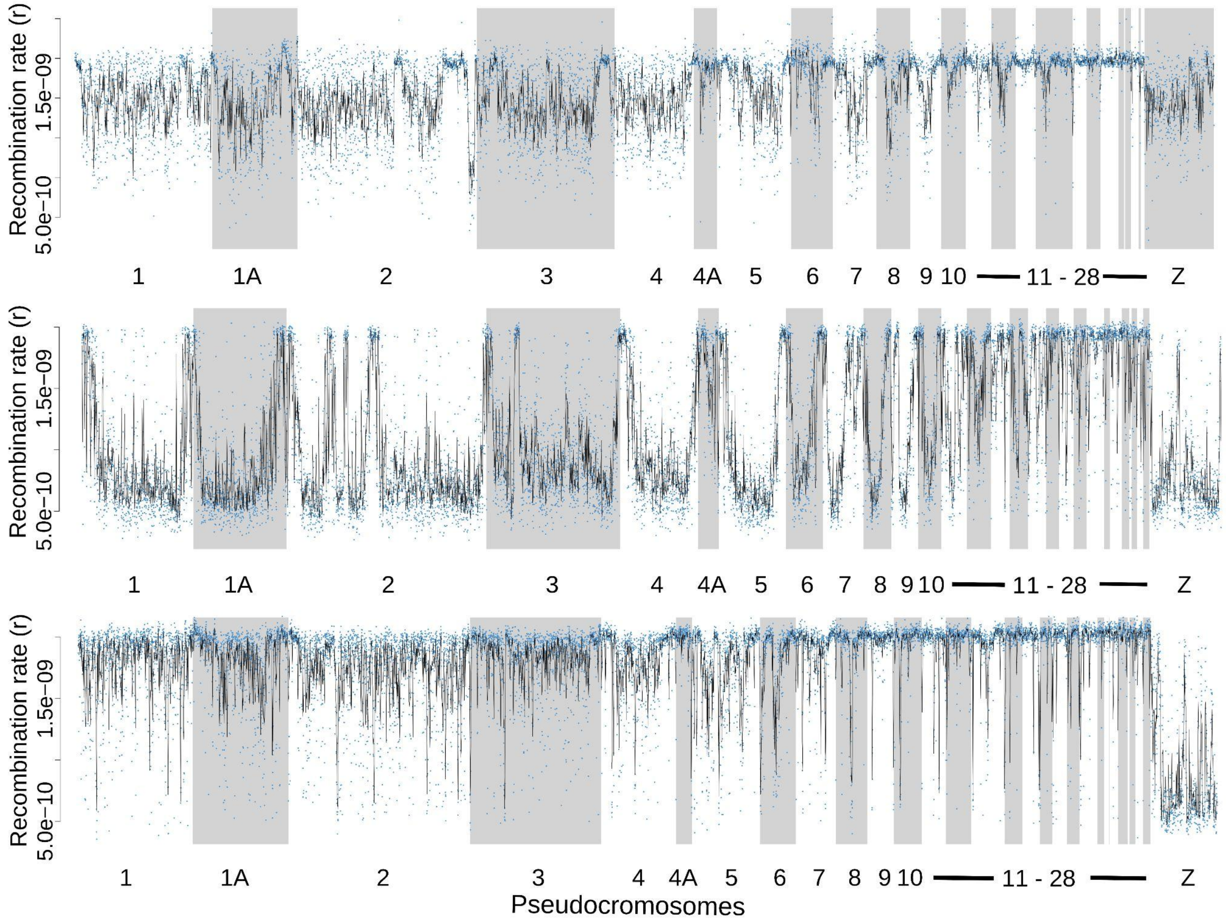


Figure S7: Recombination rate (r= recombination rate per base pair per generation) across the genome of three Amazonian species. Top - *Phlegopsis nigromaculata*; center - *Xiphorhynchus spixii*; boton - *Lipaugus vociferans*.

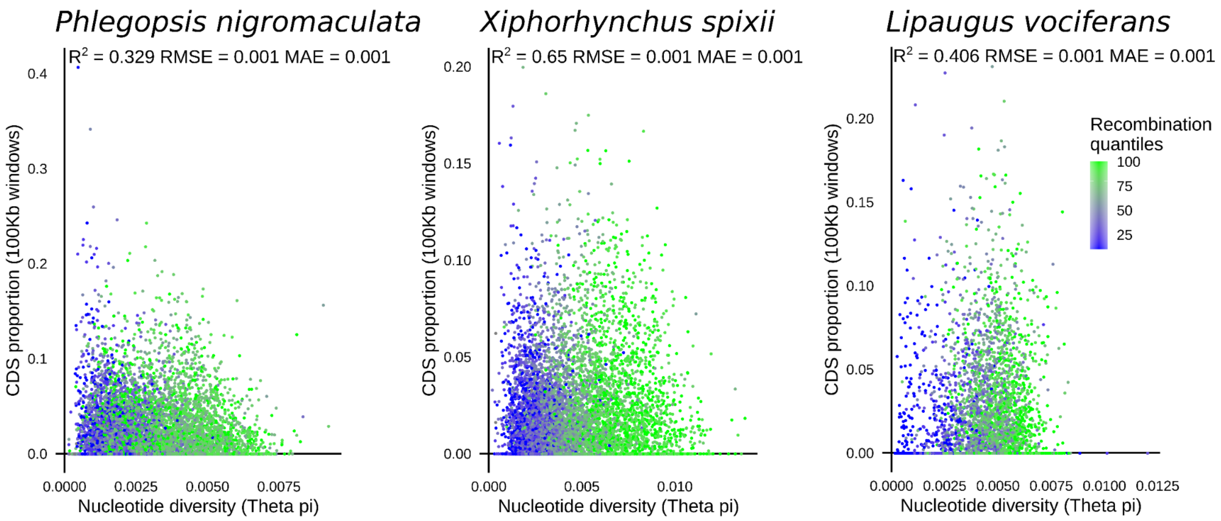

Figure S8: Relationship between nucleotide diversity, gene density and recombination rate for three species of Amazonian Birds. $R^2$, root mean square error (RMSE), and mean absolute error (MAE) were obtained with a LOESS model with nucleotide diversity as a response variable, and recombination rate and gene density as predictors.

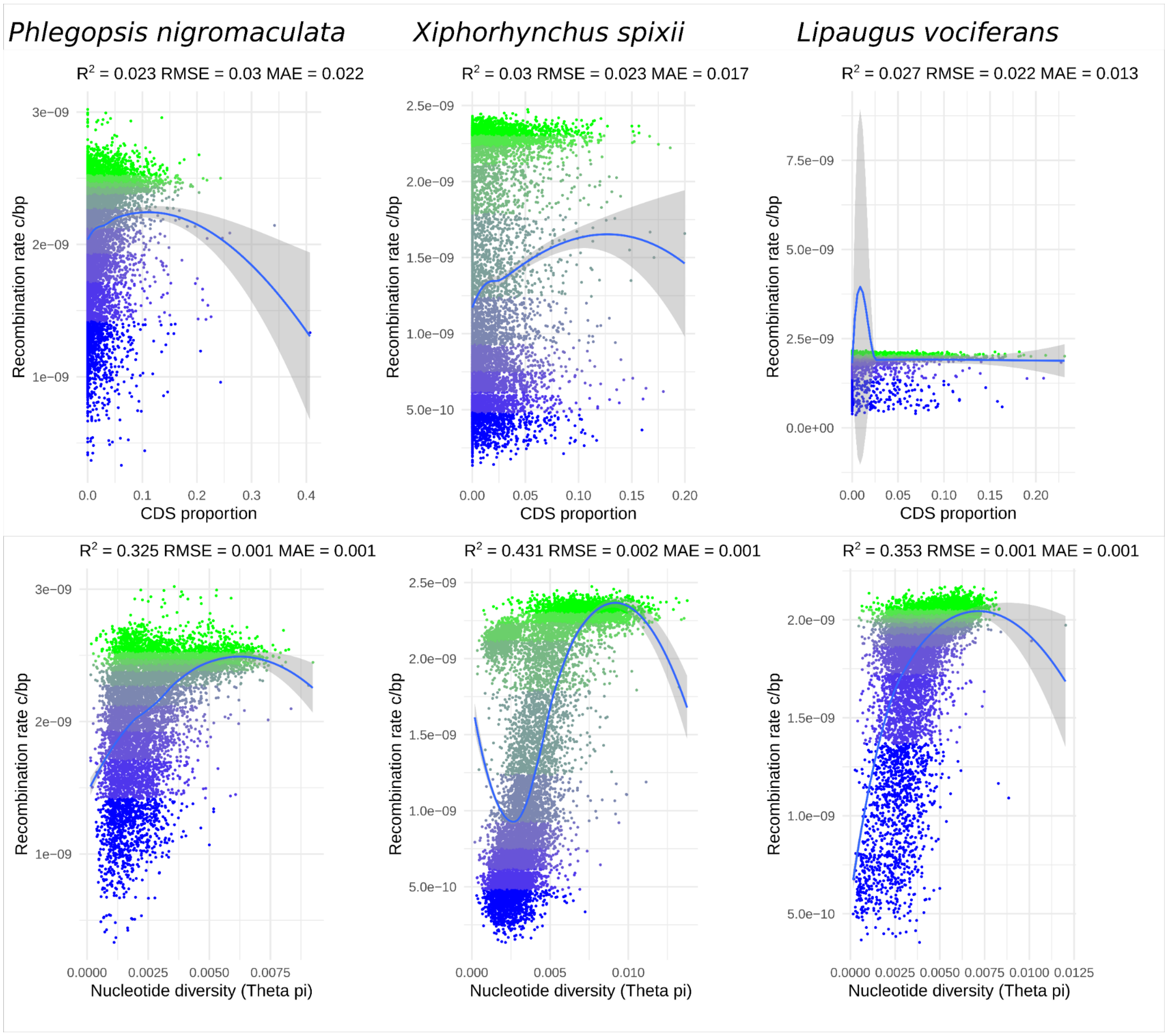


Figure S9: Relationship between nucleotide diversity, gene density and recombination rate for three species of Amazonian Birds. $R^2$, root mean square error (RMSE), and mean absolute error (MAE) were obtained with a LOESS model with nucleotide diversity or gene density as response variable, and recombination rate as predictor.

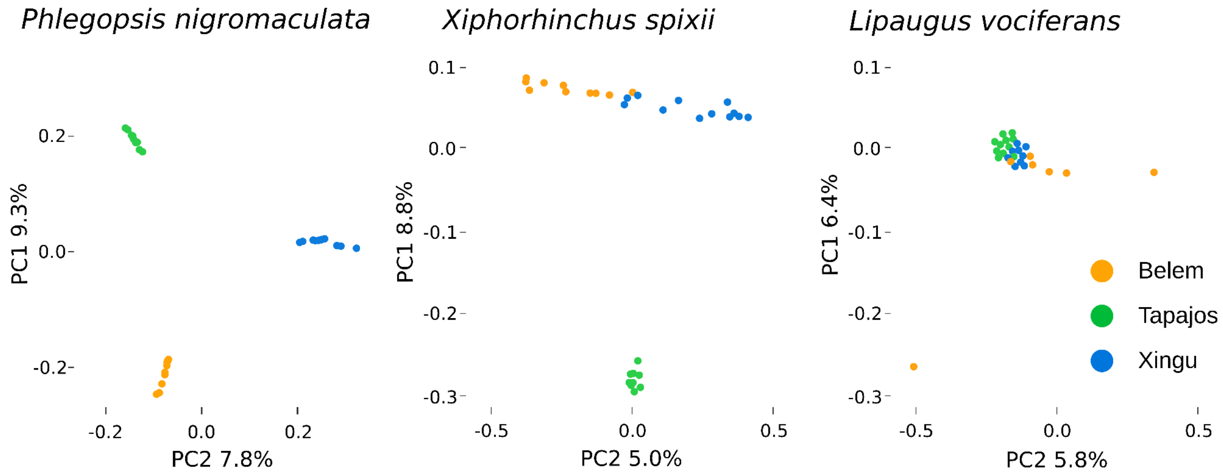


Figure S10: Principal Component Analysis for the three studied species. Colors represent samples collected in different areas of endemism, Tapajos, Xingu, and Belem.

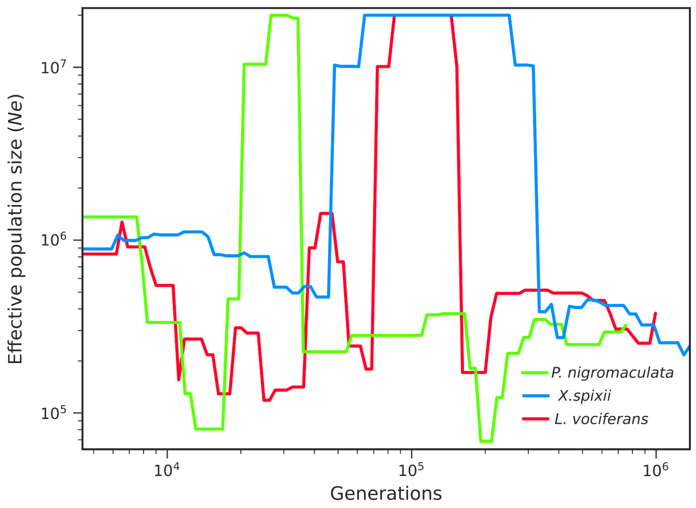


Figure S11: SMC++ estimates of effective population size through time for the Tapajos population of the three studied species. We considered a mutation rate of 2.42 x 10-9 mutations per generation and one year of generation time (Jarvis et al. 2014; Zhang et al. 2014).

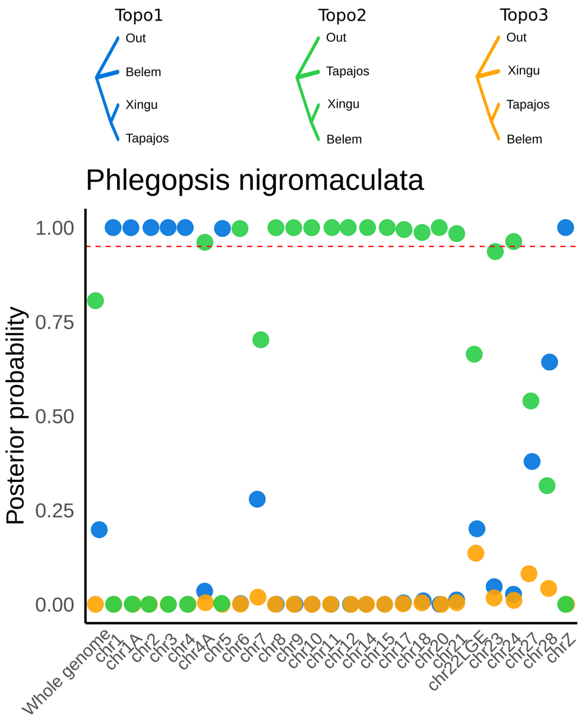


Figure S12: Species tree support across chromosomes of *Phlegopsis nigromaculata.*

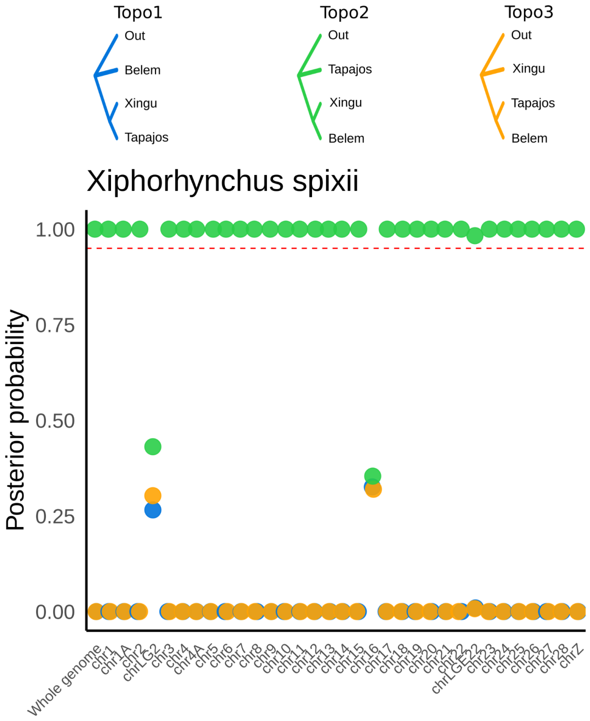

Figure S13: Species tree support across chromosomes of *Xiphorhynchus spixii*.

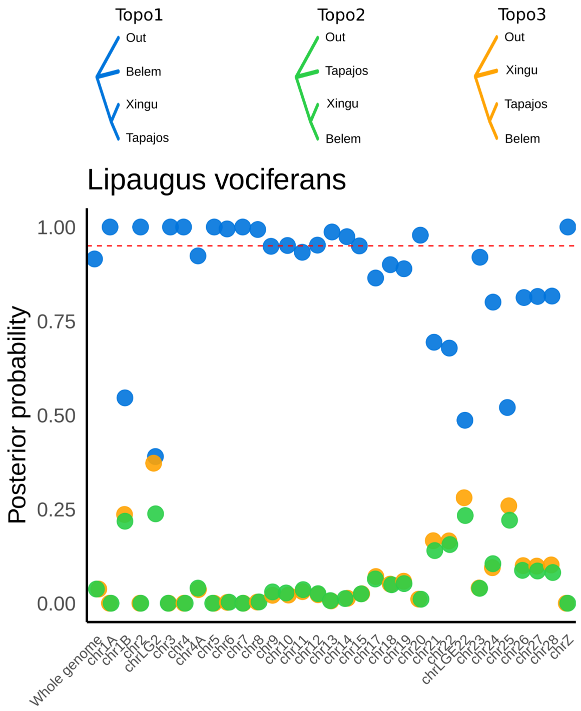


Figure S14: Species tree support across chromosomes of *Lipaugus vociferans*.

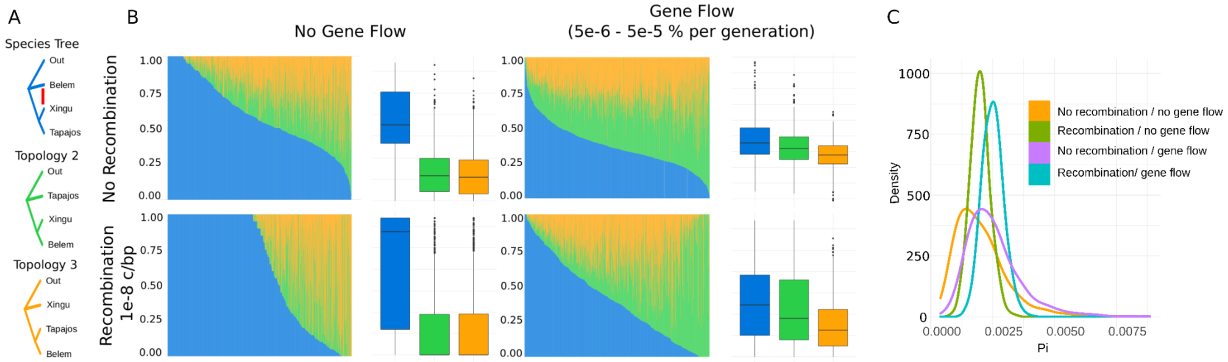


Figure S15: Topology weight and genetic diversity for four simulated models varying the presence of intra-locus recombination and gene flow between non sister lineages. Simulations were conducted assuming the species tree topology **(A)**. **(B)** For each model, the left plot shows the topology weight for the three alternative topologies for 1,000 simulations, and the right plot shows the average weight across simulations. **(C)** Distribution of nucleotide diversity across simulations for models with and without gene flow and intralocus recombination rate.

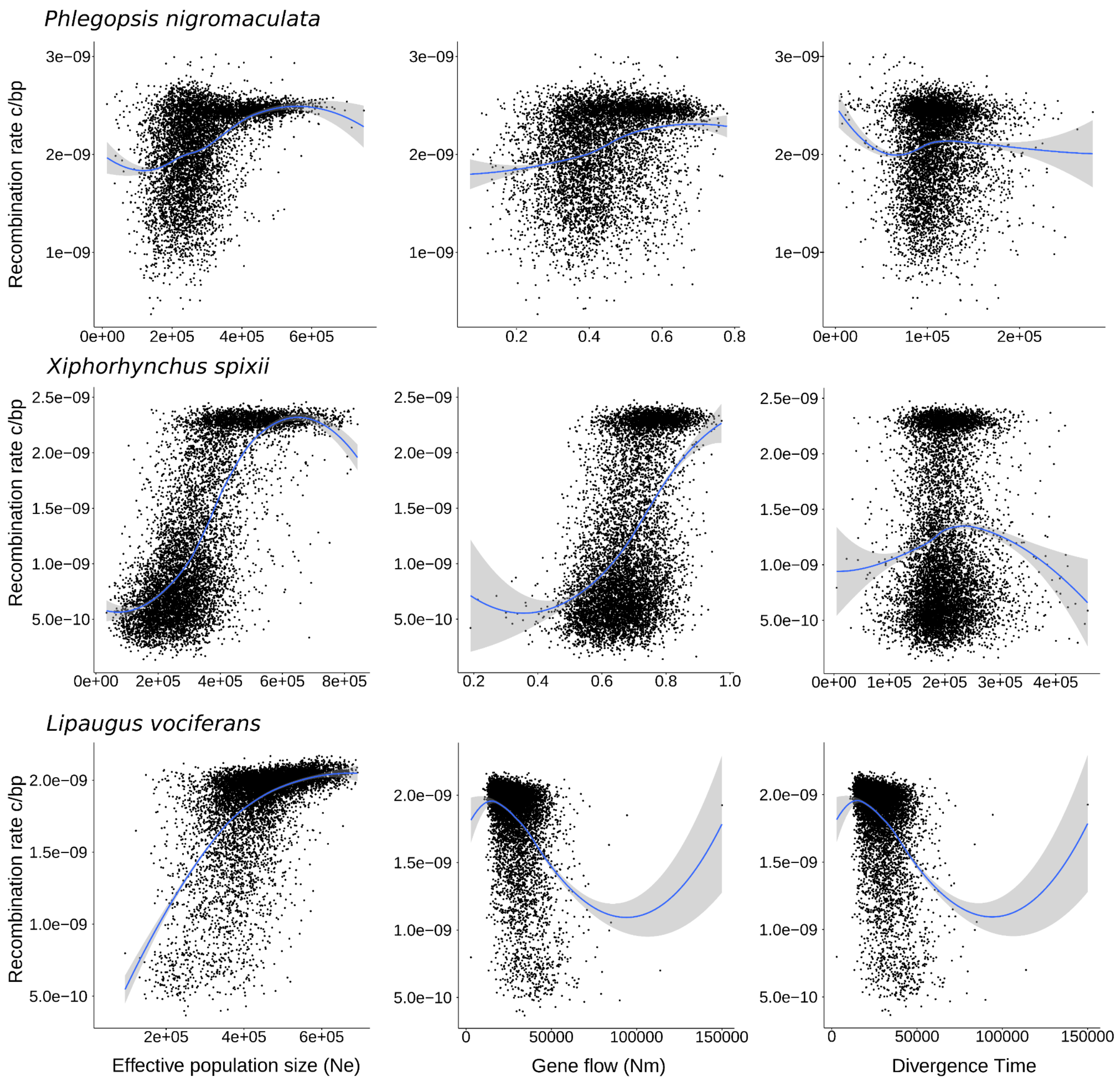


Figure S16: Association between demographic parameters estimated and recombination rate for the three studied species. Ne - Effective population size estimated for the Tapajos population (same population we used to estimate recombination rate). TMRCA - Time to the most recent common ancestor of the most recent divergence event; Gene flow - Gene flow rate between Tapajos and Xingu populations. $R^2$, root mean square error (RMSE), and mean absolute error (MAE) for each model are available in table S18.
